# *SIRT5* variants from patients with mitochondrial disease are associated with reduced SIRT5 stability and activity, but not with neuropathology

**DOI:** 10.1101/2023.12.06.570371

**Authors:** Taolin Yuan, Surinder Kumar, Mary Skinner, Ryan Victor-Joseph, Majd Abuaita, Jaap Keijer, Jessica Zhang, Thaddeus J. Kunkel, Yanghan Liu, Elyse M. Petrunak, Thomas L. Saunders, Andrew P. Lieberman, Jeanne A. Stuckey, Nouri Neamati, Fathiya Al-Murshedi, Majid Alfadhel, Johannes N. Spelbrink, Richard Rodenburg, Vincent C. J. de Boer, David B. Lombard

## Abstract

SIRT5 is a sirtuin deacylase that represents the major activity responsible for removal of negatively-charged lysine modifications, in the mitochondrial matrix and elsewhere in the cell. In benign cells and mouse models, under basal non-stressed conditions, the phenotypes of SIRT5 deficiency are generally quite subtle. Here, we identify two homozygous *SIRT5* variants in human patients suffering from severe mitochondrial disease. Both variants, P114T and L128V, are associated with reduced SIRT5 protein stability and impaired biochemical activity, with no evidence of neomorphic or dominant negative properties. The crystal structure of the P114T enzyme was solved and shows only subtle deviations from wild-type. Via CRISPR-Cas9, we generate a mouse model that recapitulates the human P114T mutation; homozygotes show reduced SIRT5 levels and activity, but no obvious metabolic abnormalities, neuropathology or other gross evidence of severe disease. We conclude that these human *SIRT5* variants most likely represent severe hypomorphs, and are likely not the primary pathogenic cause of the neuropathology observed in the patients.

## Introduction

Mitochondria are essential for maintaining the health of eukaryotes, through their functions as bioenergetic, biosynthetic and signaling organelles^1^. Consequently, mitochondria are vital for normal tissue function, and mitochondrial defects lead to a myriad of diseases^2-4^, which vary tremendously in clinical impact, from relatively subtle-age associated defects in single organ systems, to devasting multi-system disorders that are fatal early in life. This latter group includes Leigh Syndrome (LS), characterized by failure to thrive, psychomotor regression, and death in childhood^5^. In general, tissues with high energy requirements, such as brain, liver and heart, are more sensitive to mitochondrial defects and display severe clinical phenotypes^6^. Mitochondrial disease can be caused by defects in proteins involved in oxidative phosphorylation (OXPHOS) complex assembly, in mitochondrial structure, mtDNA maintenance, and other aspects of mitochondrial functioning^7-10^.

Mammalian sirtuins are a group of seven enzymes that target specific lysine modifications to regulate the biological properties of their protein targets and promote proteome integrity^11^. The sirtuin SIRT5 localizes primarily to the mitochondrial matrix, but is also present in the cytosol, peroxisomes, and nucleus. It primarily acts on the negatively-charged post-translational modifications (PTMs) succinylation, malonylation, and glutarylation^12^. The abundance of global succinylation, malonylation and glutarylation levels are markedly increased upon SIRT5 depletion in tissues, and most of SIRT5’s target proteins are enzymes involved in various metabolic pathways^13-18^. SIRT5 remains a somewhat enigmatic protein. SIRT5 plays major roles in specific cancer types, including melanoma^19^, pancreatic cancer^20^, breast cancer^21^, and AML^22^, in a context-specific manner. By contrast, in mice, under basal unstressed conditions, the effects of SIRT5 deficiency tend to be quite subtle^23^. SIRT5-deficient mice develop cardiac fibrosis and mildly-impaired heart contractile function with age^17^, and, under specific dietary conditions, hyperammonemia^24^. Whether SIRT5 might play other important, as-yet undiscovered roles in whole animals remains unclear.

Identification of patients with diseases associated with defects in specific genes and pathways has frequently provided new, unexpected insights into gene function and human biology. A number of polymorphisms and mutations have been identified in sirtuin genes in humans, some of which are associated with clinical phenotypes^25-28^. Prior work has linked *SIRT5* polymorphisms in humans to the rate of brain aging^29^ and overall longevity^30^, coronary artery disease and acute myocardial infarction^31^, and the risk of developing gastric cancer^32^. In this study, we describe two newly identified *SIRT5* variants in patients with LS-like pathology. We show that each of the two *SIRT5* variants results in reduced SIRT5 protein levels and impaired NAD^+^-dependent activity, driving an increase in global protein succinylation levels. We show that one of these variants, P114T, has minimal effects on the structure of SIRT5. Via CRISPR-Cas9, we generate mice with the P114T variant, and find that they have severely reduced SIRT5 protein levels and activity. However, P114T homozygous mice are grossly normal, and cells with this mutation are bioenergetically indistinguishable from wild-type controls. Moreover, the SIRT5 P114T variant was also present in a sibling that did not develop neuropathologic complications. The possible implications for SIRT5 and its role in mitochondrial disease are discussed.

## Results

### Whole exome sequencing identifies *SIRT5* variants in two patients with mitochondrial-like disease pathology

Whole exome sequencing of DNA from two unrelated patients presenting with severe mitochondrial-like pathology resulted in the identification of two predicted-pathogenic *SIRT5* variants. These patients did not have common copy number genetic variants according to CGH microarray analysis or major chromosomal aberrations that could associate these patients with other mitochondrial genetic variants. In the first patient, we identified a *SIRT5* homozygous missense variant (c. 340C>A, NM_012241.3) that led to a proline114 substitution to threonine (P114T) change in the SIRT5 protein. In the second patient, a distinct homozygous *SIRT5* missense variant was identified (c.382C>G, NM_012241.3) resulting in a leucine 128 to valine change (L128V).

Clinically, the first patient (a 7-year-old boy) presented with a profound global developmental delay, seizures, spastic quadriparesis, optic atrophy, blindness and deafness (Fig. 1a). An MRI of the brain, performed at 2 years of age, showed global cerebral and cerebellar atrophy with a marked reduction of white matter volume, as well as a delayed myelination pattern (Fig. 1b, left panel). Besides these severe physical abnormalities, echocardiography indicated normal cardiac function, and plasma and urine metabolite levels were largely non-remarkable. The patient was born to first cousin parents. He has an older healthy sister, and two sisters born after him who died in early infancy (age of 3 months) with progressive hypoalbuminemia and nephrotic range proteinuria, a phenotype distinct from patient 1. Neither sister had visual tracking nor responded to sounds right from the beginning. One of the sisters was heterozygous for the SIRT5 variant and her WES was negative. There was no DNA available from the second sister. The patient has a seven-month-old brother who is homozygous for the SIRT5 c.340C>A (p.P114T) variant. Thus far, this child has appropriate development and neurological examination, sitting unsupported, grasping, and transferring objects, and babbling.

**Figure 1.**
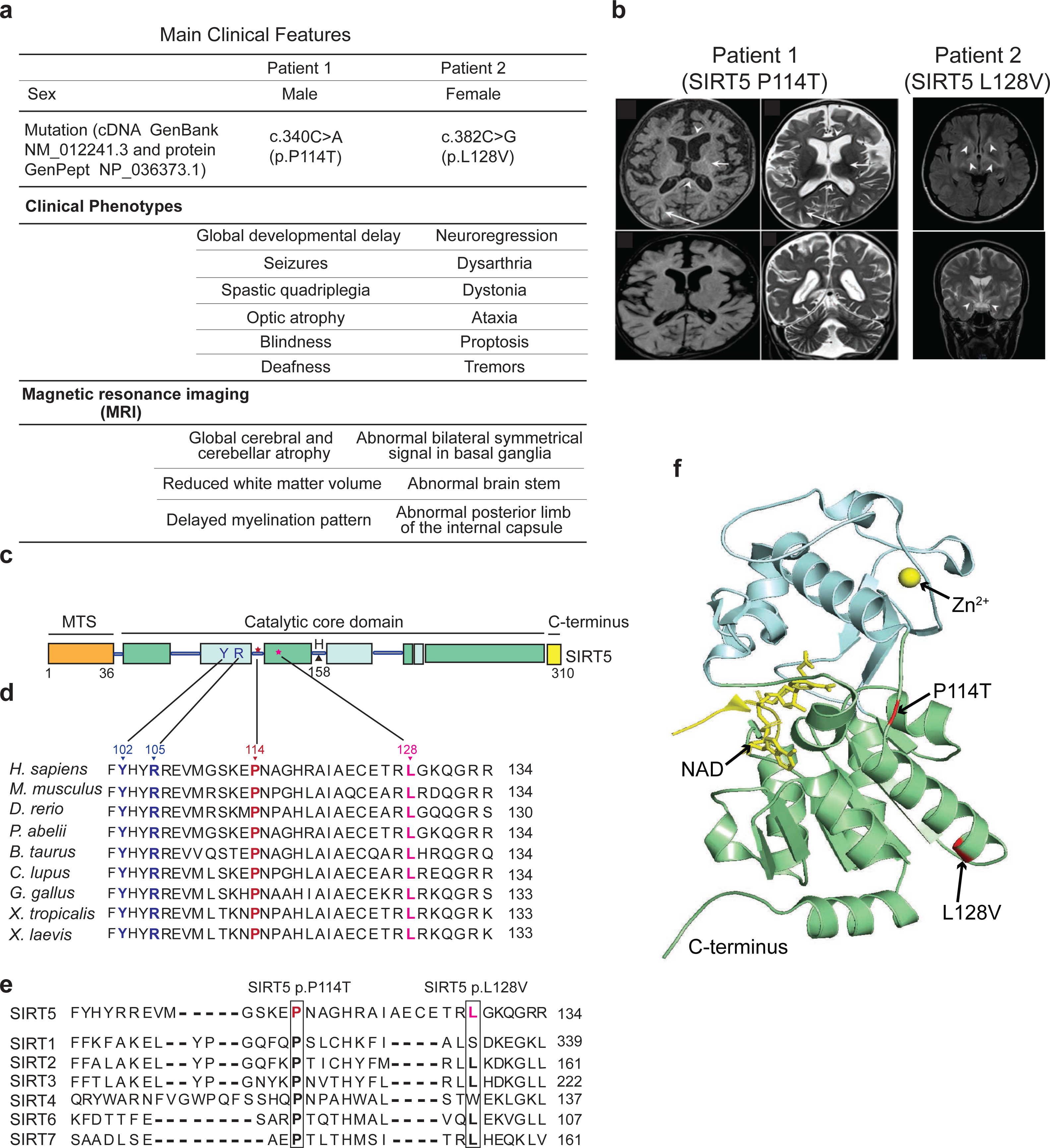
Two SIRT5 homozygous variants identified in patients with mitochondrial-like disease. (a) Clinical symptoms of the two patients. (b) MRI images of brains of the patient 1 (at the age of 2 years) and patient 2 (at the age of 25 years). (c) Schematic representation of the human SIRT5 protein structure. The catalytic core consists of Rossmann-fold (green box) and zinc-binding domains (cyan box), flanked by a mitochondrial target sequence (MTS, orange box) and the C-terminal region (yellow box). Three sites known important for catalytic activity, Y102, R105, and H158 are indicated. Two altered residues P114 (patient 1), and L128 (patient 2) are represented as star symbols. Evolutionary conservation of two mutated sites SIRT5 P114 and L128V across (d) different species, and across (e) human sirtuin family. Substrate specificity-defining residues Y102 and R105 are indicated in blue; P114 and L128 are highlighted in dark red and pink, respectively. (f) Spatial position of the altered amino acids P114 and L128 in human SIRT5. The structure was generated based on the crystal structure of human SIRT5 (PBD: 3RIY). Affected residues P114 and L128 are indicated in red. The zinc-binding domain and Rossmann-fold domain are in blue and green, respectively. The Zinc atom is included as a yellow sphere.

The second patient (a 30-year-old female) presented clinically with neuroregression, dysarthria, and dystonia (Fig. 1a). At the age of 22 years, patient 2 began to manifest ataxia and proptosis, and shortly thereafter developed tremors, that gradually resulted in loss of motor skills until she became bedridden. An MRI of the brain at age of 25 years showed bilateral symmetrical signal abnormalities in the basal ganglia, as well as in the posterior limb of the internal capsule and brain stem (Fig. 1b, right panel). Similar to patient 1, these severe clinical manifestations were not associated with plasma or urine biochemical abnormalities. This patient also manifested a borderline complex IV deficiency (229 mU/UCS versus control range 228-1032 mU/UCS) in a skeletal muscle biopsy (Supplementary Table S1, lower panel).

To gain insights into the impact of the two missense *SIRT5* variants identified in the patients, we first investigated the characteristics of the two involved residues, P114 and L128. Both P114 and L128 are located in the catalytic core domain of SIRT5 (Fig. 1c). Moreover, these two residues are highly evolutionarily conserved (Fig. 1d), as well as within the human sirtuin family (Fig. 1e). In addition, both P114 and L128 localize close to the well-established active site residues Y102 and R105 (Fig. 1d), which are critical for substrate binding^33^. Structurally, P114 lies in the flexible loop which connects the helix of the zinc-binding domain and helix α6 of the NAD^+^-binding Rossmann fold domain, and L128 is positioned in α6 helix of the NAD^+^-binding Rossmann domain (Fig. 1f). The evolutionary conservations as well as nearly coinciding positions of these two altered residues, P114 and L128, suggest their potential importance for SIRT5 function *in vivo*. Indeed, both variants P114T and L128V are predicted to be pathogenic according to Polyphen (http://genetics.bwh.harvard.edu/pph2/). Each of these variants has been detected at very low frequency in the heterozygous state in the general population (10/280,762 alleles for P114T, and 5/250,928 alleles for L128V in the Genome Aggregation Database), but neither has ever been reported in homozygous state (https://gnomad.broadinstitute.org/).

### *SIRT5* variants cause reduced levels of SIRT5 protein level and increased global succinylation levels in human fibroblasts

To understand the consequences of the SIRT5 P114T (patient 1) and L128V (patient 2) variants on SIRT5 protein and function, we generated primary skin fibroblasts from both patients and compared them to fibroblasts from healthy controls. Whole cell lysate SIRT5 protein levels were markedly reduced in patient 1 and 2 as compared to controls (Fig. 2a), despite normal *SIRT5* mRNA expression levels (Fig. 2b). To rule out the possibility that lower SIRT5 protein levels detected in patients’ fibroblasts was caused artefactually by reduced antibody affinity due to amino acid substitution, we confirmed the reduction of SIRT5 protein level using two different SIRT5 antibodies generated against distinct antigens (Supplementary Fig. S1a). In contrast to the reduced SIRT5 levels, protein levels of SIRT3 and SIRT4 remained unchanged (Fig. 2a). Moreover, complex V (ATP5A subunit) protein levels were not different between patients and controls indicating that total mitochondrial content was not affected in the SIRT5 variant cell lines (Fig. 2a).

**Figure 2.**
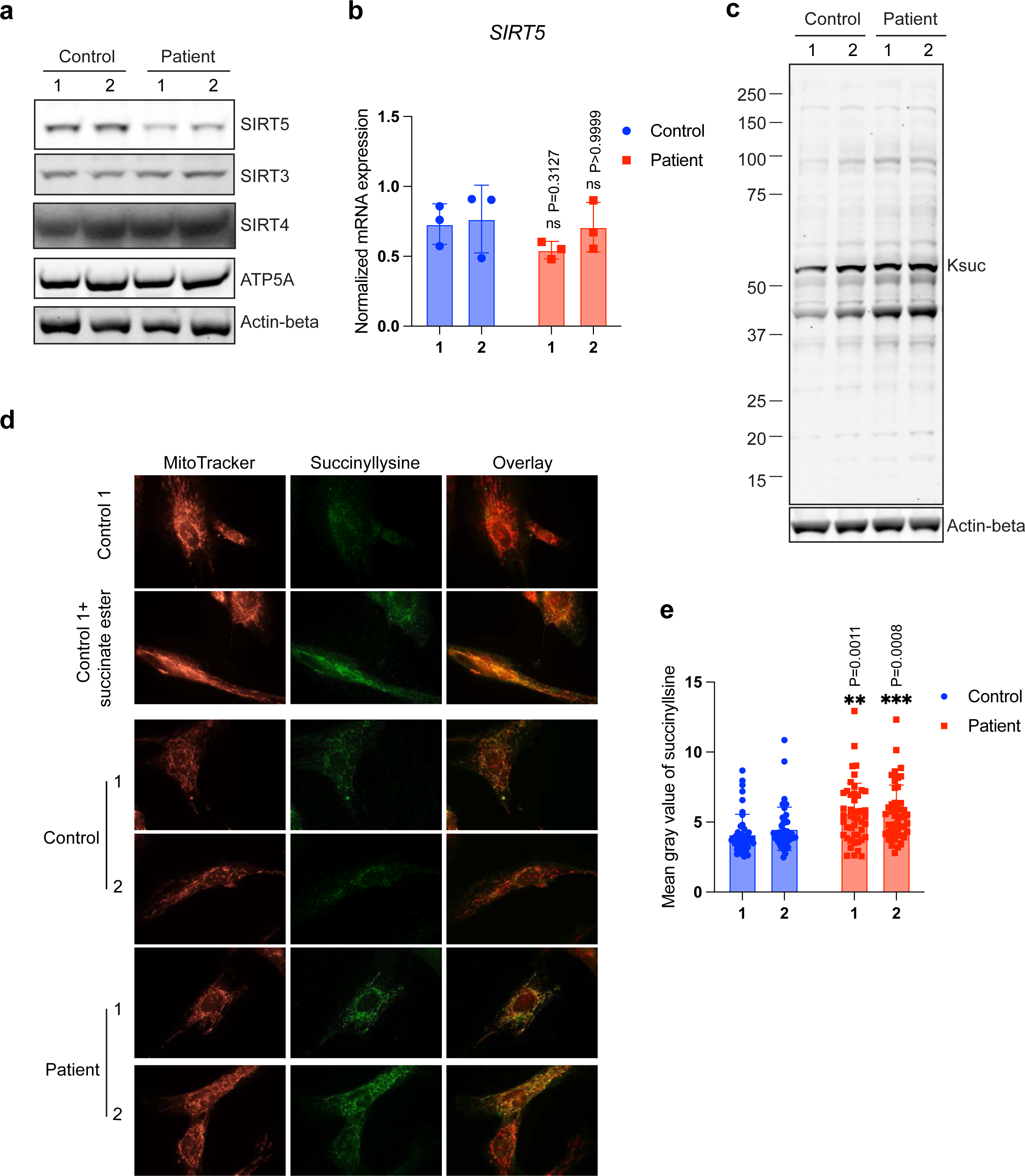
Each of the two SIRT5 variants leads to reduced SIRT5 protein level and increased global succinylation in human fibroblasts. (a) Immunoblots for SIRT5, SIRT3, and SIRT4 in whole cell lysates from control and patient fibroblasts. ATP5A and beta-actin served as loading control. (b) *SIRT5* mRNA normalized expression level in control and patient fibroblasts. Three reference genes were used for expression normalization. n = 3 independent experiments and duplicates in each experiment. (c) Immunoblots for succinyllysine and loading control beta-actin control and patient fibroblasts. (d) Representative immunofluorescence staining images of control and patient fibroblasts. Cells were stained with succinyllysine antibody and with Mitotracker orange. Control fibroblasts exposed to 4 mM cell-permeable dimethyl succinate-ester was used as positive control for succinyllysine staining. (e) Quantification of immunofluorescence staining for succinyllysine in control and patients’ fibroblasts. N = 46 images were analysed for each cell line. Data represent mean ± SD. Statistical analyses in (b) and (e) were performed using one-way ANOVA, followed by Bonferroni’s multiple comparison test. Each patient fibroblast was tested against average of controls. ** indicates P < 0.01. *** indicates P < 0.001. ns, not significant.

Next, we asked whether SIRT5 biochemical activity in cells was affected by the *SIRT5* gene variants. Because SIRT5 possesses robust desuccinylase, demalonylase and deglutarylase activities^33,34^, we first analysed these three SIRT5-related acylation levels by immunoblot. Succinyllysine levels were increased in both patient fibroblasts as compared to controls (Fig. 2c), whereas no clear difference in malonyllysine and glutaryllysine levels were detected (Supplementary Fig. S1b and S1c). Upon exposure to dimethyl succinate-ester, a cell permeable succinate analogue, we observed an increase in succinylation levels in both control and patient fibroblasts (Supplementary Fig. S1d), indicating that the succinylation machinery in the cells was not affected, and increased succinylation levels in the patient cells were due to SIRT5 dysfunction.

We tested whether localization of succinyllysine was altered in cells bearing the SIRT5 variants. Localization of intracellular succinyllysine was visualized using immunofluorescence staining. Exposure of fibroblasts to dimethyl succinate-ester confirmed that protein succinylation was most apparent in mitochondria (Fig. 2d). Consistent with immunoblot results, significantly greater succinyllysine fluorescence signal was detected in both patient fibroblasts in comparison with succinlyllysine fluorescence in controls (Fig. 2d, e). Intracellular localization of succinyllysine did not appear to be altered. Overlayed images of succinyllysine and mitotracker demonstrated that succinylation was present in both mitochondria and cytosol, especially in patient cells. Together, these results show that both *SIRT5* gene variants markedly reduced SIRT5 protein levels and significantly increased global succinyllysine levels in patient cells.

### SIRT5 variants result in loss of SIRT5 thermal stability and desuccinylase activity

To gain additional insight into the mechanism of how the *SIRT5* variants result in lower SIRT5 protein levels and higher succinylation levels, we biochemically characterized the variants *in vitro*. The patient SIRT5 variants, P114T and L128V, were engineered using site-directed mutagenesis (Fig. 3a) and these His-tagged SIRT5 recombinant proteins were purified over Ni^2+^-affinity resin purity (Supplementary Fig. S2a).

**Figure 3.**
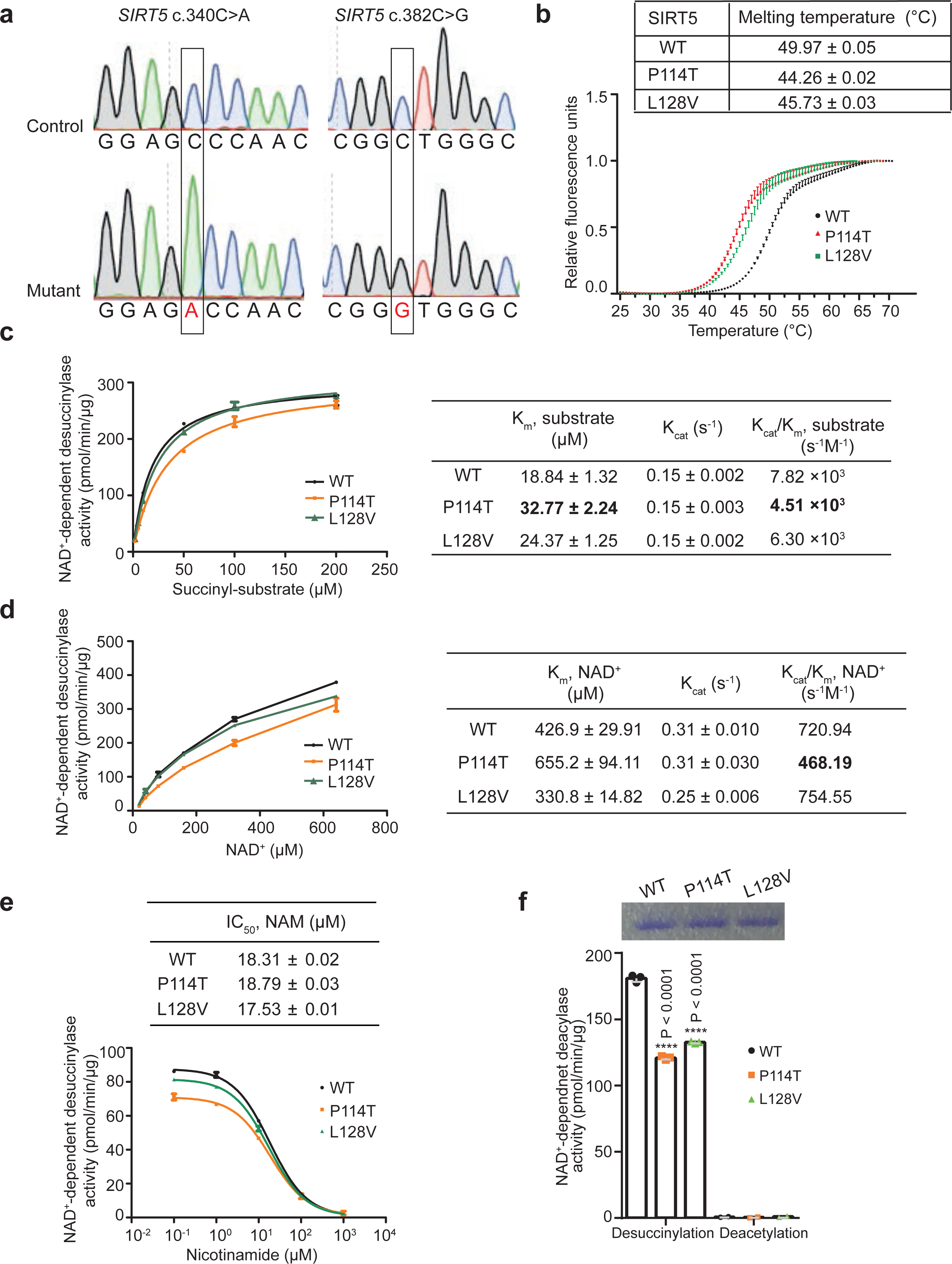
SIRT5 P114T and SIRT5 L128V variants decrease protein thermal stability and NAD+-dependent desuccinylase activity. (a) Sanger sequencing confirmed correct amino acid substitution in recombinant SIRT5; SIRT5 c.340C>A (p.P114T) represents SIRT5 variant identified in patient 1 and c.382C>G (p. L128V) represents SIRT5 variant identified in patient 2. (b) Thermal shift assay of recombinant SIRT5 WT and two variants. Triplicates for each sample. Representative for 3 independent experiments. (c) Steady-state kinetics of desuccinylation was determined by varying succinyl-peptide (2-200 μM) in the presence of 1 mM NAD+ and 0.03 μg of recombinant SIRT5 protein. Duplicates for each data point. (d) Steady-state kinetics of desuccinylation were measured by varying NAD+ (20-640 μM) in the presence of 120 μM succinyl-peptide and 0.03 μg recombinant SIRT5 protein. Duplicates for each data point. (e) IC50 values of nicotinamide (NAM) for NAD+-dependent desuccinylase activity were assessed for recombinant WT and two SIRT5 variants. Triplicates for each data point. (f) Steady-state desuccinylase and deacetylase activities were measured in recombinant WT and the two SIRT5 variants, using 10 μM acyl-peptide and 500 μM NAD+. Triplicates for desuccinylation and duplicates for deacetylation activity. Representative for 2 independent experiments. Data represent mean ± SD.

We initially assessed whether protein stability of the variants was affected. Thermal stability analysis demonstrated that the P114T as well as the L128V SIRT5 variant exhibited a lower denaturation temperature than WT SIRT5 (Fig. 3b), implying impaired thermostability, which could contribute to the reduced steady-state SIRT5 variant protein levels at physiological temperature. Next, we investigated whether enzymatic kinetics in SIRT5 variants were altered. Since succinylation levels, but not glutarylation and malonylation levels, were increased in patient fibroblasts, we focused on SIRT5’s desuccinylase activity. The Michaelis-Menten constant (K_m_) of desuccinylation of a fluor de lys succinyl-substrate by SIRT5 P114T was 74% higher, and catalytic efficiency (K_cat_/K_m_) was 42% lower, compared to WT SIRT5. For the SIRT5 L128V, the K_m_ for the succinyl-substrate was 30% higher, and catalytic efficiency was 20% lower compared to WT (Fig. 3c). Additionally, the K_m_ for NAD^+^ was 53% higher for SIRT5 P114T, and the catalytic efficiency was 35% lower compared to WT. No evident difference in catalytic efficiency with regards to NAD^+^ was observed between L128V and WT (Fig. 3d). Also, no difference was found in the IC_50_ value of the general sirtuin inhibitor, nicotinamide (NAM), for the P114T and L128V variants as compared to WT recombinant SIRT5 (Fig. 3e).

Since SIRT5 desuccinylase kinetics were only mildly decreased in the recombinant SIRT5 variants, we analysed whether desuccinylase activity was affected to a greater extent under lower substrate conditions, which could be physiologically relevant, because locally in the mitochondria NAD^+^ levels can fluctuate dramatically in response to altered metabolic conditions. Under these substrate limiting conditions, with 10 µM of succinyl-substate and a NAD^+^ concentration of 500 µM, desuccinylase activity was reduced by 30% in P114T and in L128V SIRT5 variant relative to WT (Fig. 3f). All SIRT5 alleles had neglectable deacetylation activity (Fig. 3f). In the context of limiting NAD^+^ to 10 µM, the desuccinylase activity of each of the SIRT5 patient variants was decreased further, to 50% of the activity of WT (Supplementary Fig. S2b). Together, these results indicate that the two SIRT5 variants reduced SIRT5 protein thermal stability as well as NAD^+^-dependent desuccinylase activity, likely contributing to the significant decrease in SIRT5 protein levels and increase in succinylation levels observed in the patient fibroblasts.

These studies were repeated independently in another laboratory using a different preparation of WT and SIRT5 P114T protein (Fig. S2c and d), using peptide substrates whose sequence was derived from Succinate Dehydrogenase (SDHA), a SIRT5 substrate that we and others previously identified^13,35,36^. Again, the SIRT5 P114T enzyme showed markedly reduced *in vitro* biochemical activity against succinylated, malonylated, and glutarylated substrates compared to WT control (Fig. S2e). To test whether P114T might exert dominant negative effects, we mixed WT and P114T enzymes together in an equimolar ratio. There was no apparent inhibition of WT SIRT5 by the presence of the P114T variant, suggesting that P114T did not inhibit biochemical activity of the WT enzyme (Fig. S2f). Consistent with our other results, this preparation of SIRT5 P114T, but not a catalytic H158Y mutant^13^, demonstrated reduced thermal stability compared to WT protein (Fig. S2g).

### The SIRT5 P114T mutation is not associated with gross changes in 3-dimensional structure

To further gain insight into the impaired biochemical activity of the SIRT5 P114T variant, we solved the crystal structures of human P114T and WT SIRT5 enzymes in parallel. We report the first crystal structure of human SIRT5 without a bound co-factor or substrate mimetic in the active site. The structure of WT SIRT5 was solved to 2.25 Å resolution with two protein molecules in the asymmetric unit. The apo structure of P114T was solved to 2.7 Å resolution with two molecules in the asymmetric unit. Although the apo P114T protein has a significantly lower melting temperature than the apo WT SIRT5 protein (ΔTm = -5.71 to -6.05 °C), their structures are nearly identical aligning with an overall RMSD of 0.518 Å as calculated by SMS in Coot^37^ (Fig. 4a). The cause for the lower melting temperature of the variant structure may be attributed to the fact that a proline, which is a hydrophobic residue having a very rigid structure, was mutated to a more flexible, hydrophilic threonine residue. In the WT structure, Pro114 forms ten hydrophobic interactions with the sidechains of Lys148, Ala149, His118, Leu145 and Glu113 and its carbonyl oxygen forms a hydrogen bond with Arg119 (Fig. 4b). Thr114 forms only five hydrophobic interactions with the sidechains of Arg119, His118, Ala149. Although Thr114 loses the hydrogen bond between its carbonyl oxygen and the side chain of Arg119, it gains a hydrogen bond between its hydroxyl group and the carbonyl oxygen of Leu145 (Fig. 4c). The formation of this new hydrogen bond could interfere with hydrogen bond between the backbone amides of Leu145 and Ala149 in the WT structure. We conclude that the SIRT5 P114T variant shows impaired biochemical activity and thermal stability, but no dramatic change in 3-dimensional structure as assessed by crystallography.

**Figure 4:**
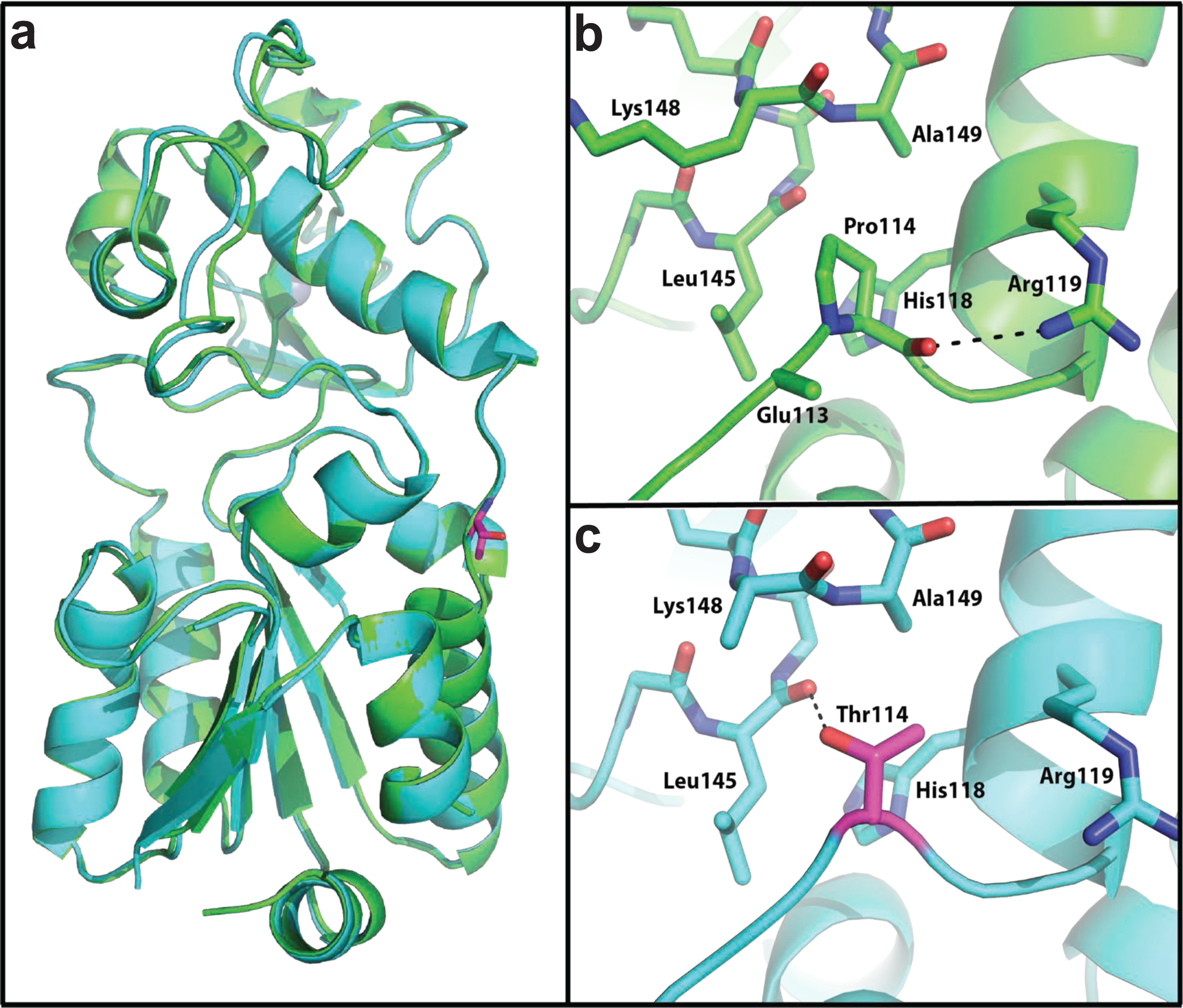
Structural alignment of the apo structures of SIRT5 WT (green) and P114T (cyan). a) Thr114 is shown as sticks with carbons colored magenta. b) Pro114 and the residues that interact with it are shown as sticks. c) Thr114 and the residues that interact with it are shown as sticks. Hydrogen bonds are depicted as dashed lines.

### P114T mice show reduced SIRT5 levels and elevated levels of SIRT5’s target PTMs

Since the biochemical activity and thermal stability of the SIRT5 P114T variant were impaired, but homozygosity of the P114T SIRT5 variant was also observed in an apparently healthy sibling of patient 1, we aimed to gain a more detailed understanding of the impact of the P114T SIRT5 variant in mice. To model the effects of P114T homozygosity in mice, we used CRISPR-Cas9 technology to generate mice with the P114T variant (Fig. 5a and Fig. S3a and b). We generated three distinct lines based on three mosaic founders, and confirmed the identity of the correct mutation using Sanger sequencing on genomic DNA derived from tail snips (Fig. 5b). Heterozygous and homozygous P114T animals were born at the expected Mendelian ratios and were grossly unremarkable (Fig. 5c and Fig. S3c). Overall body weight, as well as heart and brain weight, were indistinguishable between WT controls in P114T homozygous adults (Fig. 5d).

**Figure 5:**
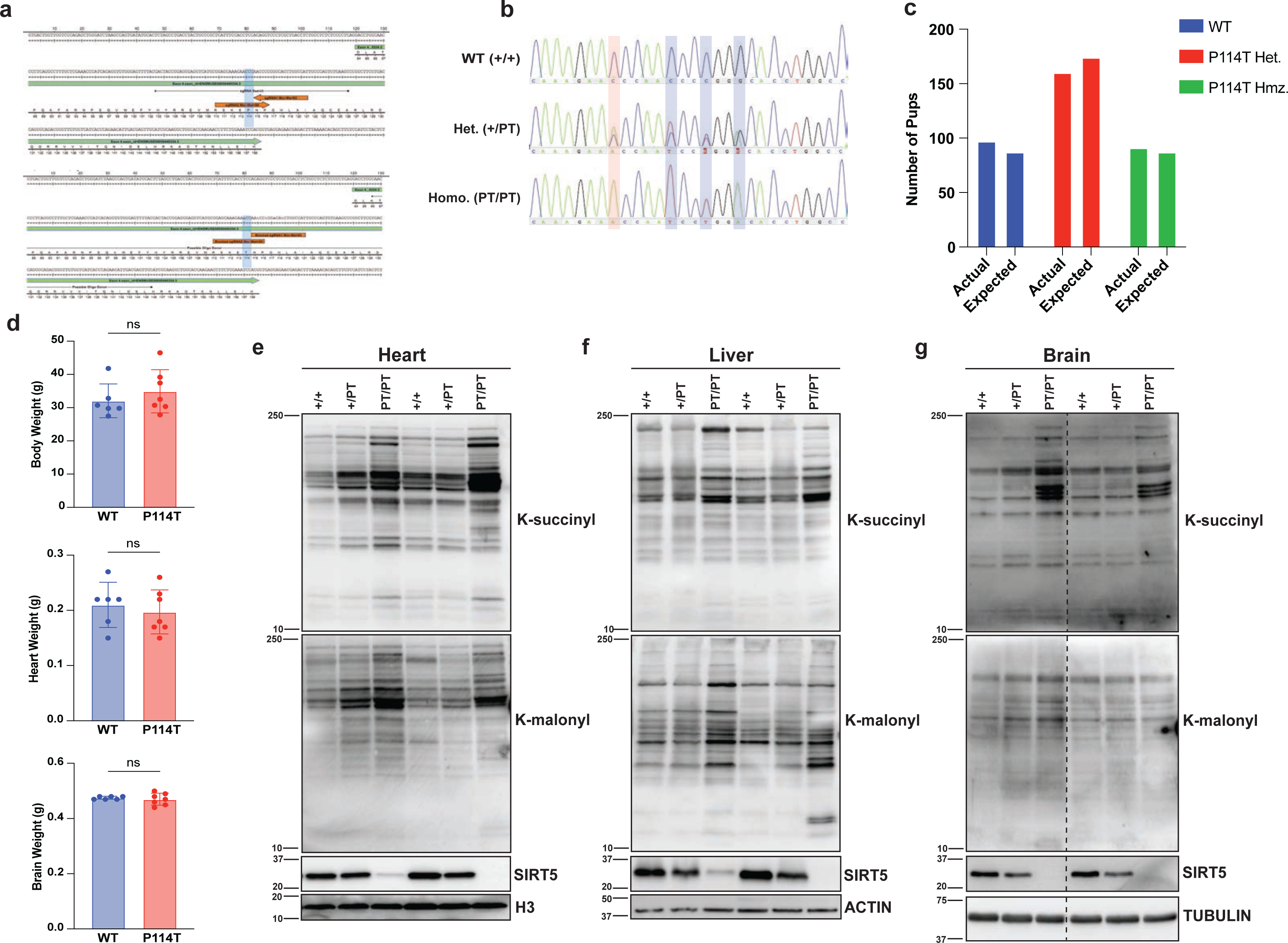
P114T variant leads to reduced SIRT5 protein levels and increased global lysine succinylation and lysine malonylation levels in mouse tissues. a) Mouse *Sirt5* exon 4 showing guide RNA target site for CRISPR/Cas9 to introduce the c.340C>A (p.P114T) change. b) Representative Sanger sequencing peaks of genomic DNA derived from tail snips of *SIRT5* WT, P114T heterozygous and P114T homozygous mice. c) P114T pups were born at the expected Mendelian ratios (ns, Chi-squared test). d) WT and P114T homozygous mice show no difference in body weight, heart weight and brain weight. e-g) Immunoblots showing reduced levels of SIRT5 protein and increased Ksucc and Kmal in P114T heterozygous and P114T homozygous tissues (heart, liver, and brain) compared to WT controls.

Given the reduced thermal stability of the P114T variant, we assessed SIRT5 protein levels in tissues of mice with this mutation. SIRT5 protein levels were markedly reduced in P114T heterozygotes, and virtually undetectable in homozygotes (Fig. 5e-g). We interrogated levels of SIRT5 target PTMs in tissues from P114T heterozygotes and homozygotes. Consistent with prior findings in *Sirt5* null animals, Ksucc and Kmal levels were markedly elevated in P114T variant hearts, liver, and brain (Fig. 5e-g). Kglu levels appeared to be elevated as well, but the relatively poor quality of the antibody reagent available made it difficult to render firm conclusions regarding changes in levels of this mark (Fig. S3d and e).

Similar to the results from tissues, SIRT5 protein levels were markedly reduced in P114T heterozygous mouse embryonic fibroblasts (MEFs), and virtually undetectable in homozygous MEFs (Fig. 6a). To assess whether variant SIRT5 might be lost via proteasomal-mediated degradation, we incubated WT, P114T heterozygous, or variant cell lines in the presence or absence of the proteasomal inhibitor MG132 (Fig. 6b). As a control, we also probed for MYC, a well-known proteasomal target^38^. We were unable to restore cellular SIRT5 levels in P114T cells with MG132, although induction of MYC protein by MG132 was robust in all three genotypes (Fig. 6b). To further investigate the impact of the P114T mutation on SIRT5 levels, we measured *Sirt5* mRNA levels in MEF lines of all three genotypes. Unexpectedly, and discrepant with our results from patient cells, we found that *Sirt5* mRNA levels were reduced by roughly half in P114T heterozygotes, and further decreased in homozygous cells (Fig. S4a). The molecular basis for this reduction in *Sirt5* mRNA levels in the P114T variant is unclear. We explore possible implications of this finding in the Discussion section.

**Figure 6:**
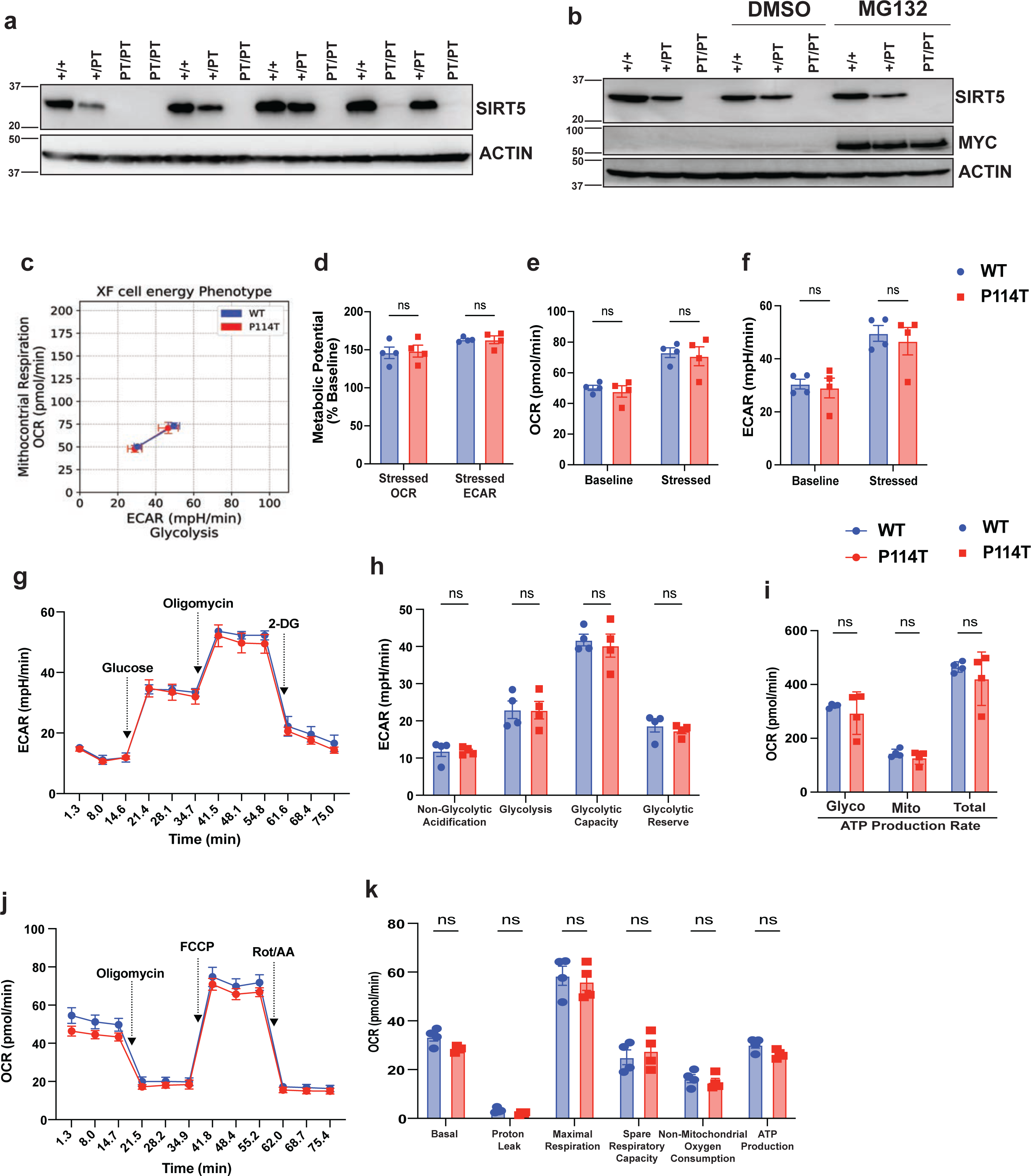
The P114T mutation has no apparent impact on the bioenergetics of mouse embryonic fibroblasts (MEFs). a) Immunoblot showing reduced levels of SIRT5 in P114T homozygous MEFs. b) Treatment with the proteasomal inhibitor did not rescue reduced levels of SIRT5 protein in P114T MEFs. c-f) Cellular bioenergetic phenotype of P114T MEFs is indistinguishable from WT. P114T MEFs maintain glycolytic function (g-h), glycolytic, mitochondrial, and total ATP production rates (i), and mitochondrial respiration (j-k) compared with littermate WT MEFs. Mitochondrial respiration, glycolytic stress tests, and ATP production rates were measured using a Seahorse XFe96 Analyzer. All rates are normalized to total protein content per sample. Each point represents an individual biological sample (average of n=6). Error bars represent SEM. Significance calculated by t-test. OCR, oxygen consumption rate; ECAR, extracellular acidification rate. Each point represents an individual biological sample (average of n=3). Error bars represent SEM. Significance calculated by t-test.

### P114T variant fibroblasts are bioenergetically unremarkable

Given the strong clinical phenotypes observed in patients with homozygous *SIRT5* variants, we subjected P114T MEFs and controls to detailed metabolic analysis using the Agilent Seahorse Analyzer. First, we performed the XF cell energy phenotypic test to measure the oxygen consumption rate (OCR), a measure of mitochondrial respiration, and the extracellular acidification rate (ECAR), a measure of glycolysis, under baseline and stressed conditions (Fig. 6c-6f). Relative to WT littermate controls, SIRT5 P114T MEFs did not show any changes in OCR and ECAR under basal conditions (baseline, in presence of nonlimiting substrate quantity) or under induced energy demanding condition (stressed, in presence of stressor compounds). Furthermore, both WT and P114T MEFs exhibit similar metabolic potential (calculated as percentage increase of stressed OCR over baseline OCR, and stressed ECAR over baseline ECAR), a measure of cellular ability to meet an energy demand via respiration and glycolysis. Likewise, both glycolytic and mitochondrial ATP production rates were unaffected by the P114T mutation (Fig. 6i). We also performed the glycolysis stress test (Fig. 6g and h), and the mitochondrial stress test (in the presence of either glucose or galactose) (Fig. 6j and k, and S. Fig. S4b and c) to measure other parameters associated with glycolysis or mitochondrial respiration. No significant differences were observed between WT and P114T MEFs. We conclude that the P114T mutation does not confer any striking respiratory or glycolytic phenotypes in MEFs under the conditions tested.

### The P114T mutation does not affect brain anatomy in mice

Owing to the neuropathology identified in the patients with biallelic *SIRT5* variants, we performed detailed neuroanatomic analysis on the P114T mice. Midline sagittal brain sections of littermate WT and P114T homozygous mice at 21-50 weeks revealed no abnormalities. Brain weights were indistinguishable between WT controls and P114T homozygotes (Fig. 5d). No evidence of neuronal loss in hippocampus, cerebellum, or cortex was detected by immunofluorescence staining for NeuN (Fig. S5a-f). Similarly, no evidence of astrocytic or microglial activation was detected by staining for GFAP or IBA1, respectively (Fig. 7a-k and Fig. S5a-f).

**Figure 7:**
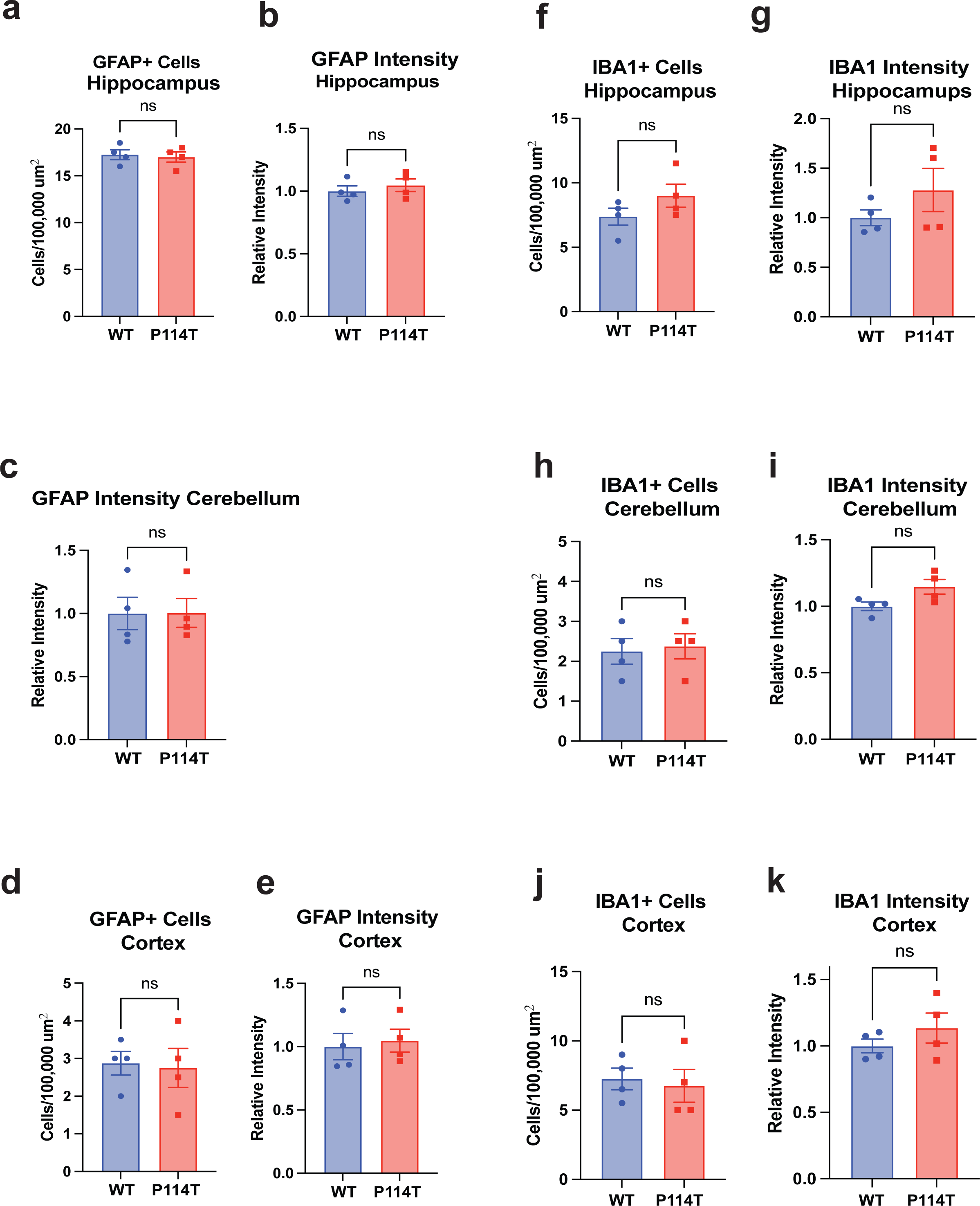
P114T mice do not show evidence of gliosis. Brains from littermate WT and P114T mice were stained for GFAP (Fig. 7a-e and Fig. S5a-c) and IBA1 (Fig. 7f-k and Fig. S5d-f) and co-stained for NeuN. Images were taken from the hippocampus (Fig. S5a and d), cerebellum (Fig. S5b and e), and cortex (Fig. S5c and f). The number of cells staining positive for GFAP and IBA1 and the intensity of GFAP (Fig. 7a-e) and IBA1 (Fig. 7f-k) staining were quantified for each image. Data are mean ± SEM. N = 4 for all groups. n.s., not significant. Student’s paired t-test (df = 6, t = (A) 0.333, 0.72; (B) 0.03; (C) 0.21, 0.34; (D) 1.47, 1.21; (E) 0.28, 2.32; (F) 0.35, 1.09.

## Discussion

In the present study, we describe two novel homozygous SIRT5 variants in two unrelated patients with mitochondrial disease-like pathologies. Comprehensive biochemical and molecular analyses revealed that each of the two SIRT5 loss-of-function variants led to a reduction in SIRT5 thermostability and protein levels, along with reduced biochemical activity, coupled with significantly higher succinylation levels in patient-derived fibroblasts. We solved the structure of one of these variants, P114T, and identified only minimal changes from the WT structure. P114T homozygous cells are bioenergetically intact, and mice with this change show no marked abnormalities, including in the CNS. Notably, the absence of a strong phenotype in P114T mice is highly reminiscent of the effects of SIRT5 knockout (KO) in mice^12^. Given the constellation of findings presented here, we believe that these SIRT5 variant alleles most likely represent severe hypomorphs, rather than dominant negatives or gain-of-function mutations in both human and mice.

Roles for SIRT5 in metabolic health and biology have been supported in previous studies by evidence generated in cells and genetically engineered mouse models, revealing a failure of SIRT5 KO mice to respond appropriately to stress, e.g. cardiac pressure overload. However, discrepancies in the biological impacts of SIRT5 deficiency have been observed among different studies, rendering SIRT5’s key biological functions in benign tissues difficult to elucidate. Our observations that human SIRT5 genetic variants are associated with dramatic clinical phenotypes emphasizes the direct relevance of SIRT5 for human health. A key issue raised by our studies is why there is an apparent discrepancy between humans and mice regarding the effects of SIRT5 loss of function. This may reflect an important underlying difference in SIRT5 biology between the two organisms. Likewise, it may be that compensatory mechanisms to overcome any putative deleterious effects of SIRT5 loss-of-function are more effectively recruited in mice than in humans. It also may be that the phenotypic effects of SIRT5 deficiency are highly dependent on modifier loci present in the genetic background, in humans and perhaps in mice. The very long lifespan of humans versus mice may offer another explanation; perhaps the short lifespan of experimental mice provides an insufficient window of time to permit sufficient accumulation of hyper-succinylated and -malonylated proteins to induce pathology. In this regard, a recent study demonstrated that conditional deletion of *Sirt5* in cardiomyocytes induced only slow subsequent accumulation of modified proteins in the heart, which occurred over the course of many weeks^39^. Alternatively, SIRT5 has been shown to regulate the urea cycle, and to inhibit blood ammonia accumulation in response to fasting and other dietary interventions^24^. Perhaps the sequelae of hypothetical episodic hyperammonemia induced by SIRT5 loss-of-function are more evident in humans than in mice, in light of our much longer lifespan. Militating against this possibility, mouse models of urea cycle deficiency generally closely phenocopy the associated human diseases^40^.

One interesting finding of these studies is that *Sirt5* mRNA levels are severely reduced in mouse cells bearing the P114T mutation, but not in humans. Currently, we do not know why this is the case. *SIRT5* mRNA expression is positively regulated by PGC-1IZ, and inhibited by AMPK^41^. It is possible that SIRT5 might regulate one or both of these factors, in an auto-regulatory loop, thereby effectively modulating its own expression, in a species-specific fashion. In this regard, multiple reports indicate that deficiency of SIRT3, the primary mitochondrial deacetylase, affects AMPK and PGC-1IZ levels and activity^42^. Whatever the underlying mechanism may be, it is likely that reduced steady-state *Sirt5* mRNA levels in the P114T variant contribute to the markedly reduced SIRT5 protein levels in this background in the mouse, in addition to the inherent reduced stability of the P114T protein.

In summary, in this study we have identified variants in *SIRT5* associated with severe mitochondrial disease, but are likely not pathogenic. These variants are associated with reduced SIRT5 levels and activity, albeit (for P114T) little change to SIRT5’s gross 3-dimensional structure. Mice with this alteration show accumulation of succinylated and malonylated proteins highly reminiscent of *Sirt5* knockout animals. These mice lack the severe phenotypes found in the two human patients. Taken together, our cellular and mouse physiologic data on the two new SIRT5 alleles indicate that these are likely not pathogenic variants, at least on their own. However, SIRT5, like many other sirtuins, might function as a modifier protein that, in combination with other genetic or environmental stressors, could influence physiology^12^. Thus, future studies might uncover more subtle phenotypes in the P114T mice similar to those observed in the *Sirt5* knockout strain – for example, defects in the urea cycle^24^, age-associated cardiac fibrosis^17^ and/or cardiac stress sensitivity^35,43^.

### Limitations of the study

The small number of pedigrees and patients characterized in this work renders it challenging to arrive at definitive conclusions regarding the impact of hypomorphic variants in *SIRT5.* It is possible that more detailed biochemical analysis, for example using unbiased metabolomic studies, might uncover consistent defects conferred by *SIRT5* loss-of-function, in the human or the mouse. Likewise, specific stress conditions, such as fasting, altered diet, or even aging, might uncover novel phenotypes in the *Sirt5* P114T strain.

## Methods

### Patient ethics

Written and oral informed consent for diagnostic and research studies was obtained for both subjects in accordance with the Declaration of Helsinki and following the regulations of the local medical ethics committee.

### Cell Culture

Human fibroblasts from skin biopsies of healthy subjects and two patients with *SIRT5* variants were used in this study. Control fibroblasts included: #13370, #MW28, #14308, and #14321; patients fibroblasts harbouring *SIRT5* variants: #13395 and #13706. All cell culture media supplements were obtained from Gibco, Thermo Fisher Scientific, unless otherwise stated. Human primary skin fibroblasts were routinely cultured in fibroblasts full growth medium (medium 199 (#M3769, Sigma-Aldrich, which is formulated with 5.5 mM glucose) supplemented with 10% foetal bovine serum (#06Q3501K), 2 mM Glut-Max (#35050038), and 1x antibiotic-antimycotic mix (100 IU/ml penicillin/streptomycin, and 25 µg/mL Amphotericin B, #15240062)). Cells were incubated at 37°C and were maintained in an atmosphere containing 5% CO_2_. Cells were passaged once every 4-5 days when reaching 90% confluency by trypsinization (#15400054) for 2 min at 37°C. All fibroblast cell lines were tested mycoplasma-free (MycoSensor QPCR assay, #302107, Agilent Technologies).

### Western blotting

After harvest by trypsinization and centrifugation (5 min, 300 x *g*, 4°C), the cell pellet was lysed in lysis buffer (50 mM Tris/HCl pH 7.4, 2 mM nicotinamide, 1 µM trichostatin A and protease inhibitor cocktail (#04693159001, Roche)), then sonicated on ice. Protein concentration was determined with BioRad DC assay (#5000116), and equalized to the same level with lysis buffer, after which it was incubated with NuPAGE LDS Sample Buffer (# NP0007, Invitrogen) and 50 mM dithiothreitol at 95°C for 5 min. Protein samples were separated on a 4-12% Bis-Tris Gel (#NW04125BOX, Invitrogen) and electroblotted onto nitrocellulose membrane (#88018, Invitrogen). The membrane was then probed with anti-SIRT5, anti-SIRT3, anti-SIRT4, anti-succinyllysine, anti-glutaryllysine, anti-malonyllysine, anti-complex V, anti-actin beta or anti-histone3, followed by appropriate secondary antibody (IRDye 800CW or HRP-linked IgG). Images were obtained using the Odyssey or Bio-Rad ChemiDoc XRS Systems. Actin beta and histone H3 were used as loading controls. Antibodies information is listed in Table 1. Uncropped blots are provided as supplementary information.

**Table 1.**
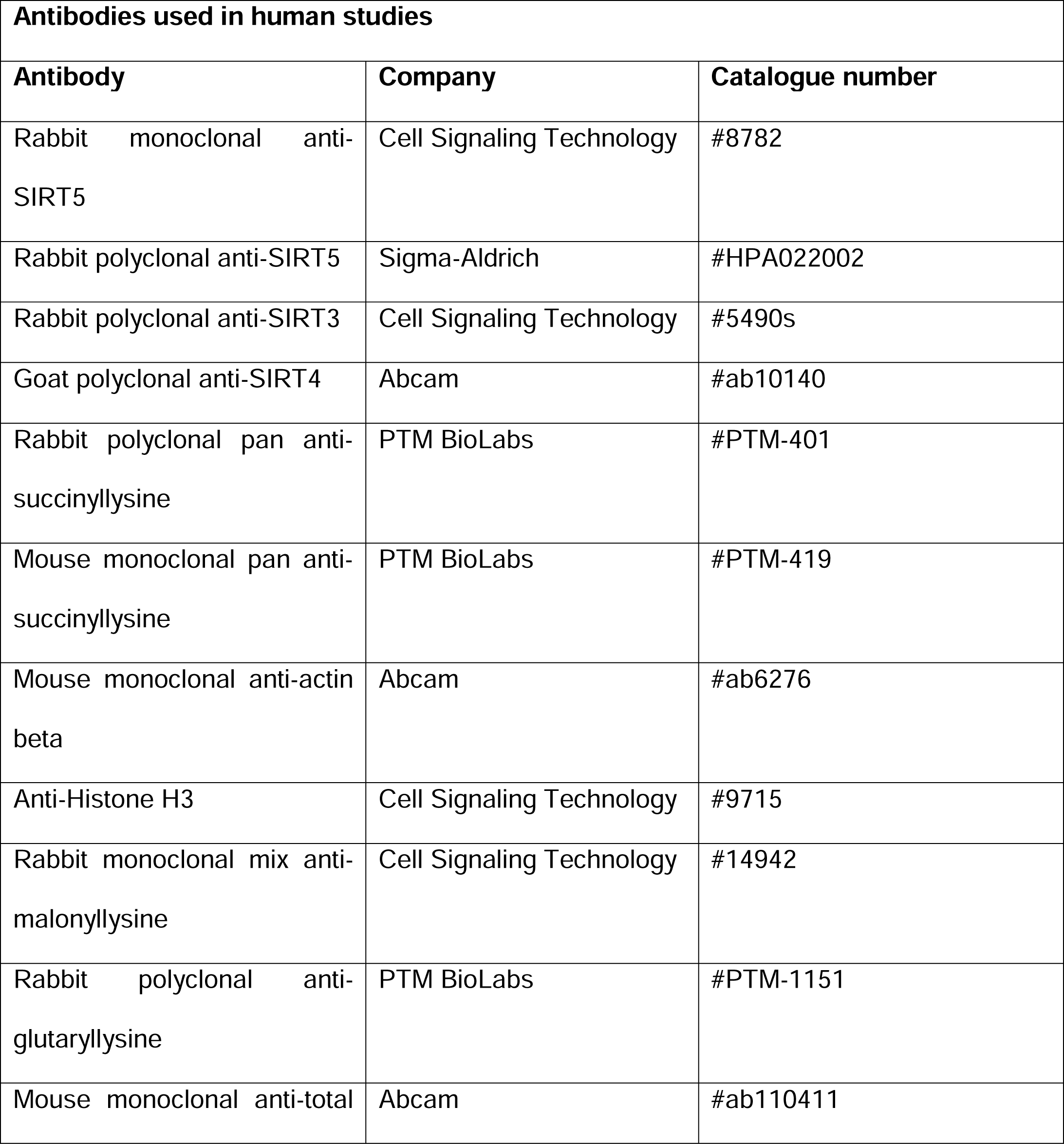
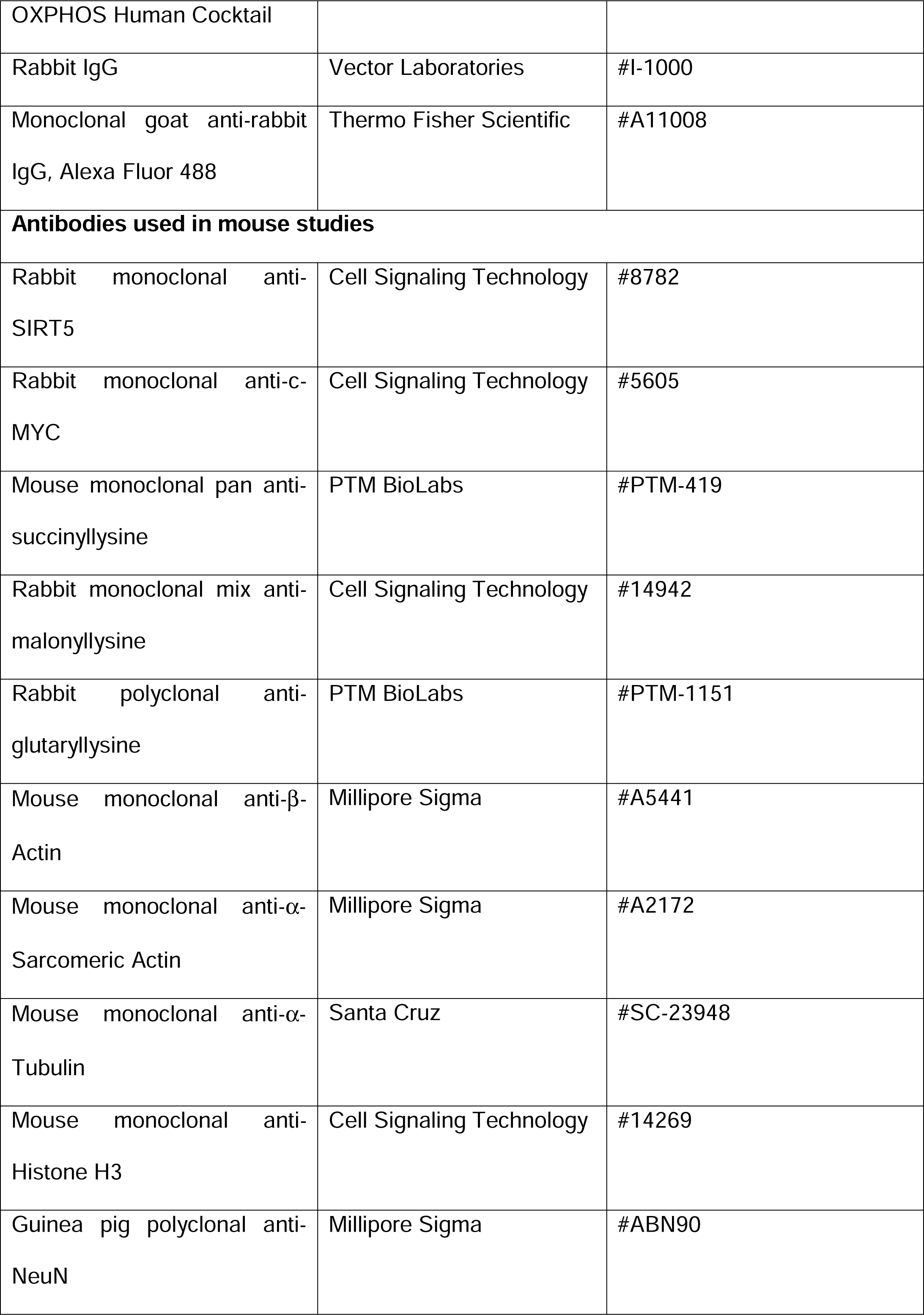
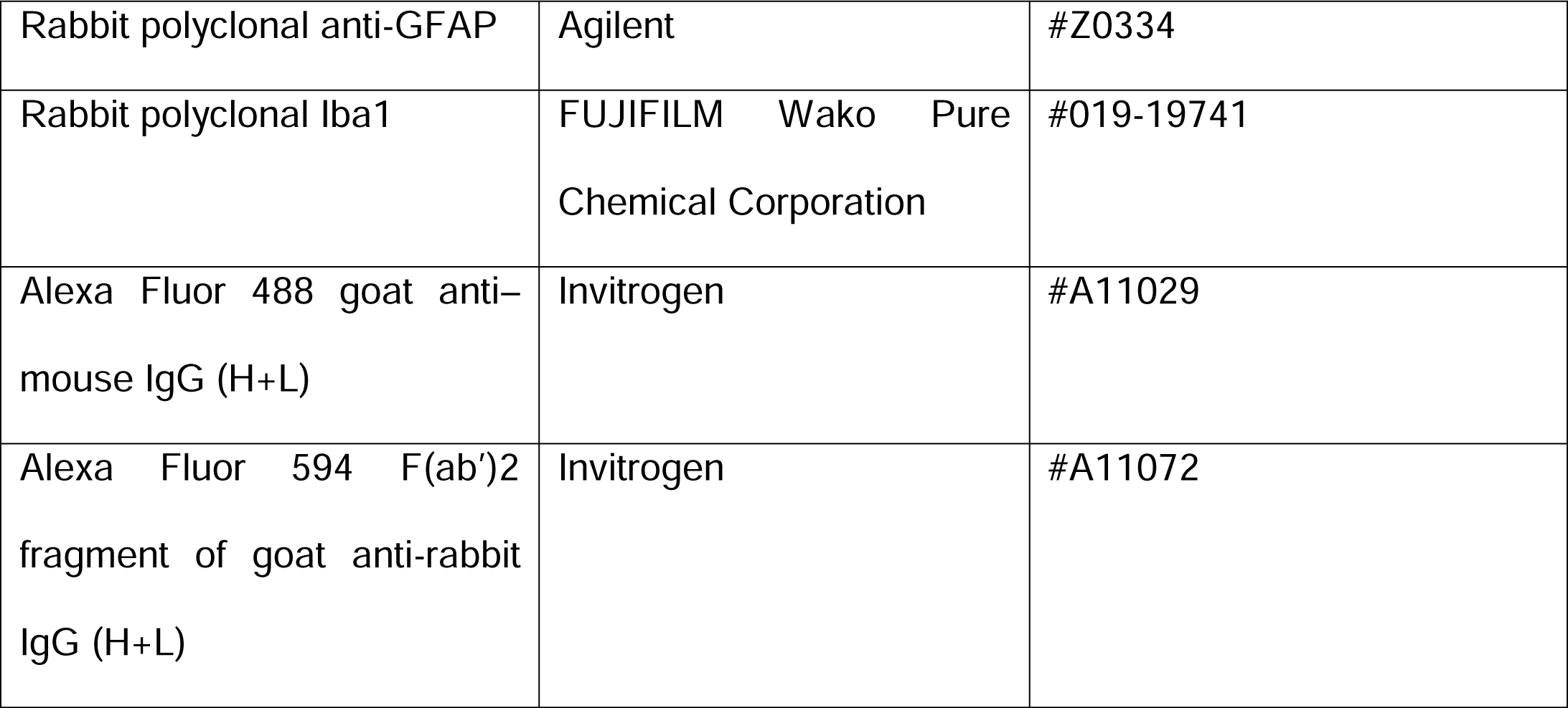
List of antibodies.

### Immunofluorescence

Fibroblasts were seeded on sterile 24 mm cover glasses in 6-well plate. Cells were cultured in fibroblast complete growth medium as mentioned before and allowed to grow for 48 hrs before the assay. For succinate exposure, cells were cultured in fibroblasts full growth medium supplemented with 4 mM dimethyl succinate-ester (#112755). Mitochondria were visualized with 400 nM MitoTracker Orange (#M7510, Molecular Probes) diluted in medium 199. Cells were fixed with 4% formaldehyde solution (#252549) at 37°C for 20 min, then were permeabilized by 0.2% Triton X-100 in Tris-buffered saline (TBS). Cells underwent blocking (2% BSA in TBS, 2% goat serum, 0.1% Triton X-100, 0.05% Tween 20). Then, they were incubated with either pan anti-succinyllysine or rabbit IgG at room temperature for 90 min, washed and incubated with the secondary AlexaFluor488 conjugated goat anti-rabbit antibody at room temperature for 60 min. Finally, cells were counterstained with 1 µg/ml of 4′,6-diamidino-2-phenylindole (DAPI, #D9564), mounted with Fluoromount G (#0100-01, SouthernBiotech). Images were taken by Leica DM6b upright microscope and quantification was performed by ImageJ 1.52. 46 images of each cell lines were quantified. All chemicals used in immunofluorescence experiments were from Sigma-Aldrich, unless otherwise stated.

### Mutagenesis of SIRT5 plasmid

His-SIRT5 expression plasmid was obtained from Addgene (plasmid #25487). Site-directed mutagenesis of WT SIRT5 was performed using the QuickChange Lightning site-directed mutagenesis kit (#210518, Agilent Technologies), according to the manufacturer’s instruction. Plasmids were amplified in competent *E.coli* and purified by using an Endofree plasmid max kit (#12362, Qiagen). Plasmids DNA were sent for Sanger sequencing to confirm the introduced *SIRT5* variants (BaseClear, the Netherlands). Mutagenesis primers are listed in Table 2.

**Table 2.**
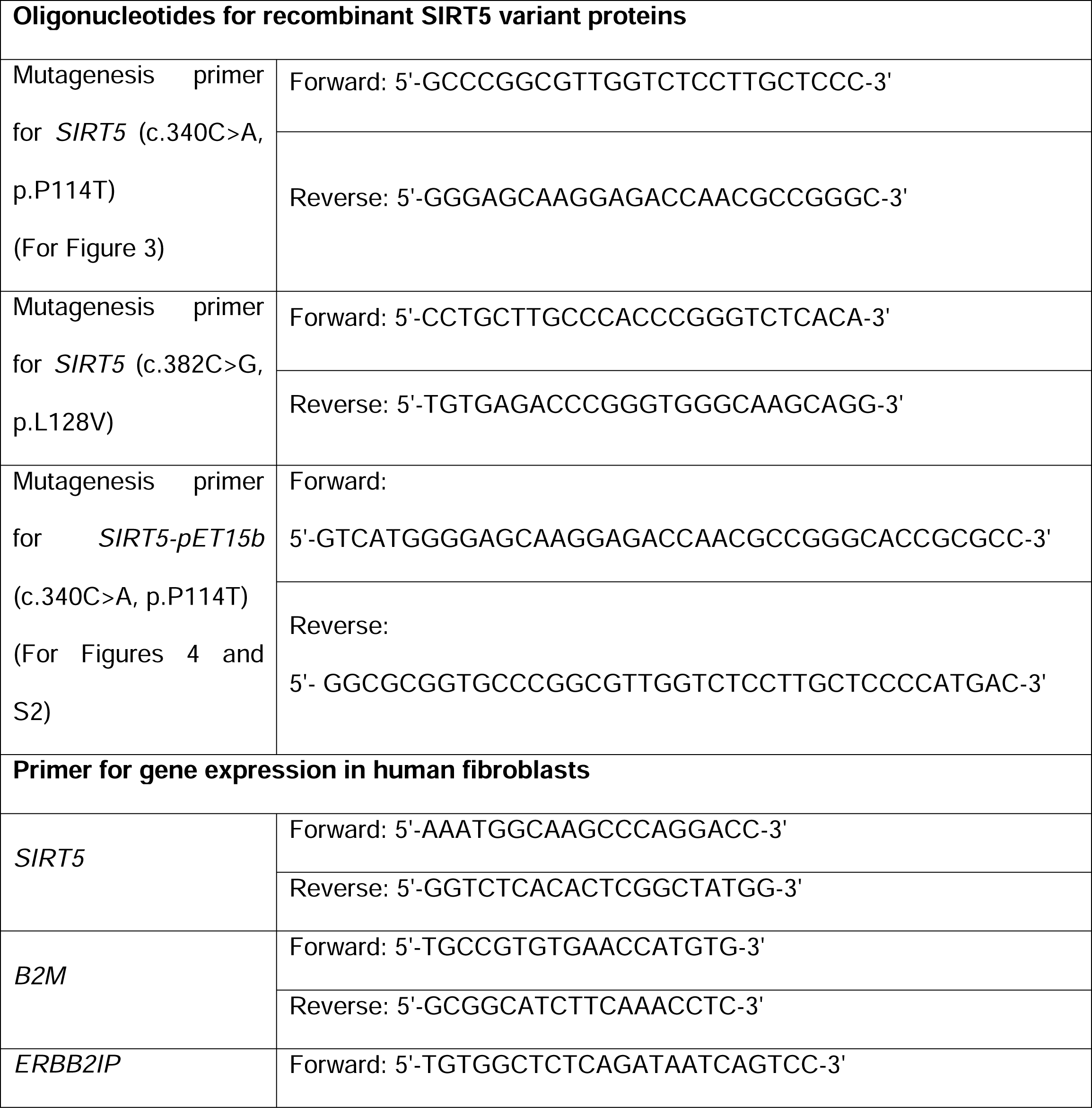
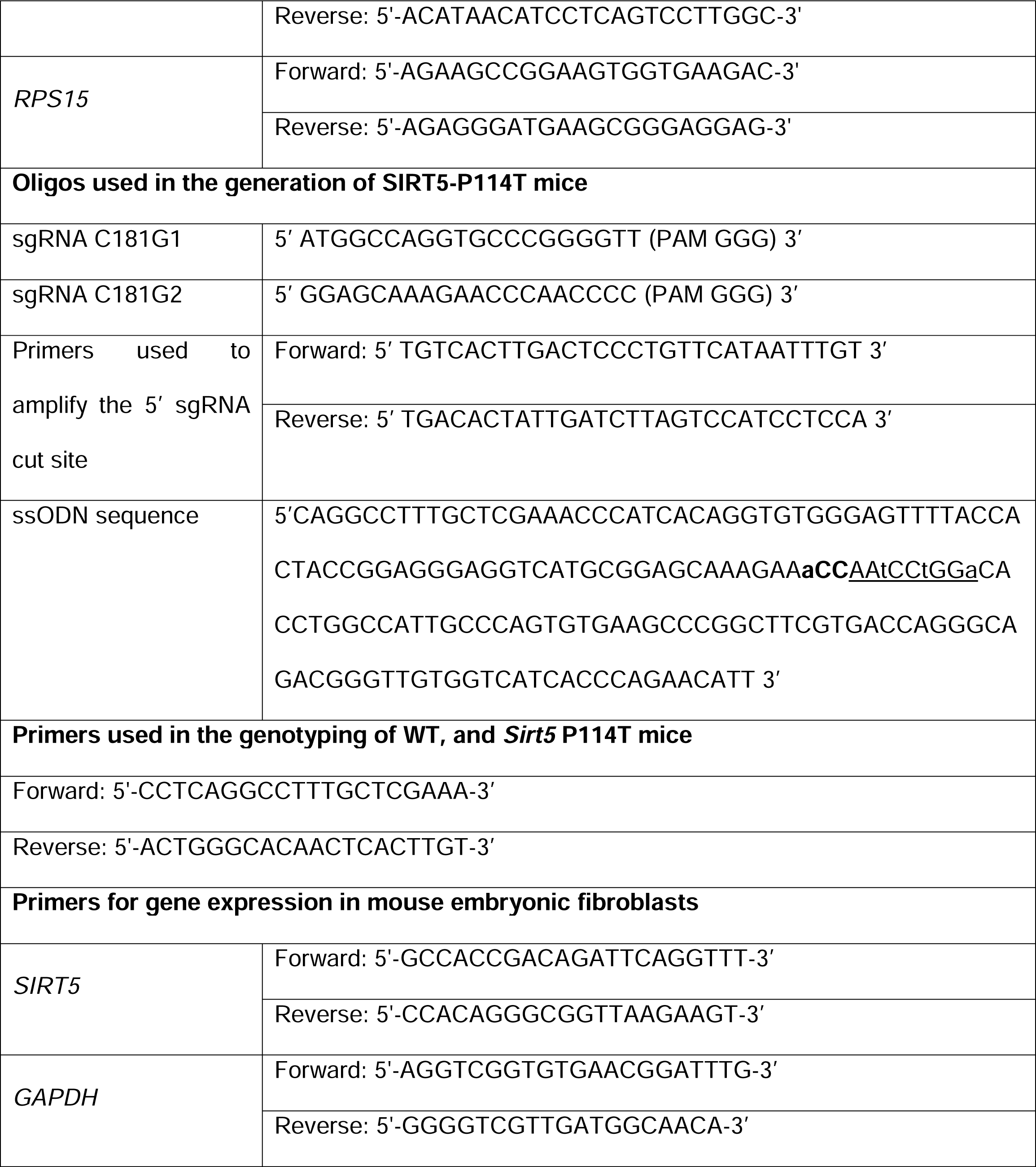
List of primers/oligos.

### Expression and Purification of Recombinant His-SIRT5 (Figure 3)

His-tagged WT and variant SIRT5 proteins were expressed in the BL21(DE3) *Escherichia coli* (*E.coli*) strain. The transformed BL21DE3 cultures were grown in Luria broth (LB) medium containing kanamycin (50 µg/ml) at 37°C until OD=0.25 was reached, after which the culture was allowed to cool down to 22°C. SIRT5 expression was induced with 500 mM isopropyl β-d-1-thiogalactopyranoside (IPTG) and the cells were harvested 18 hrs after the induction by centrifugation at 6000 x *g* for 15 min at 4°C, and snap frozen at -80°C. To purify His-SIRT5, the *E.coli* pellet was resuspended in 10 mM Tris/HCl pH 8.0, 500 mM NaCl supplemented with 1 mg/ml lysozyme, and sonicated on ice. After centrifugation at 12,000 x *g* for 15 min at 4°C, SIRT5 WT and variants were purified by Ni^2+^-affinity resin (#V8823, Promega), and eluted with 25 mM Tris/HCl pH 8.0, 100 mM NaCl, and 500 mM imidazole. Prior to use in further experiments, imidazole was removed from the eluted His-SIRT5 using ultra-0.5 centrifugal filter units (UFC501096, Merck Millipore). 500 µl of the recombinant SIRT5 elution fractionation was added to the ultra-0.5 centrifugal filter units and centrifuged at 14,000 x *g* for 15 min at room temperature. Flow-through was discarded and 450 µl of exchange buffer (25 mM Tris/HCl pH 8.0, 100 mM NaCl) was added, and the centrifugation step was repeated. Washing was repeated twice. Protein concentration of recombinant SIRT5 was subsequently determined by Bradford assay (#23200, Thermo Fisher Scientific).

### SIRT5 enzymatic assay and steady-state kinetic analyses

Desuccinylase activity of recombinant His-SIRT5 WT, P114T and L128V variants was measured with the fluor de lys SIRT5 fluorometric drug discovery assay kit (#BML-AK5130001, Enzo). To assess the Michaelis-Menten constant (K_m_) of SIRT5 for succinyl-peptides, steady-state rates were measured by varying succinyl-substrate (2-200 µM) in the presence of 1 mM NAD^+^. For the determination of K ^+^, activity was measured under a range of NAD^+^ (20-320 µM) and 120 µM of succinyl-substrate. The enzymatic reaction took place at 37°C water bath with shaking (60 rpm) for 10 min. The reaction was stopped after 10 min by incubating with 1x developer and 2 mM nicotinamide at 25°C for 15 min. The fluorescence was read in half-area black 96-well plate (#3686, Corning) with excitation at 360 nm and emission at 460 nm on a Biotek Synergy HT microplate reader. Fluorescence signal of negative control (presence of succinyl-substrate and absence of NAD^+^ in the reaction system) was subtracted from each experimental sample. For analysis of His-SIRT5 sensitivity to NAM, fixed concentration of succinyl-substrate (10 µM) and NAD^+^ (500 µM) and varying concentration of NAM (0.1 -1000 µM) were present in the reaction system. For analysis of His-SIRT5 desuccinylase activity under limiting conditions, succinyl-peptide was fixed at 10 µM, and NAD^+^ input was used as indicated in the figure legends. 0.03 µg of freshly prepared His-SIRT5 (WT, P114T, and L128V SIRT5) were used in all the experiments. For kinetics parameters calculation, data were fitted to the Michaelis-Menten equation in GraphPad Prism software (version 5.04). For IC_50_ of NAM for each His-SIRT5, data were fitted to the log(inhibitor) vs response (three parameters) equation in GraphPad Prism software (version 5.04).

### Protein thermal shift assay (Figure 3b)

Wild-type and two variants of His-SIRT5 were freshly prepared as mentioned in “Expression and Purification of Recombinant His-SIRT5” and used in the experiment. After protein concentration determination, all His-SIRT5 were equalized to 320 µg/ml with exchange buffer (25 mM Tris/HCl pH 8.0, 100 mM NaCl) containing 20x Sypro Orange (#S5692, Sigma-Aldrich, delivered at 5000x). Samples (25 µl) were added to qPCR plate and analysed with Bio-Rad CFX96 Real-Time System C1000 thermal cycler. The temperature was increased at a rate of 0.5°C/min ranging from 25-95°C, and fluorescence signal was monitored in the FRET channel of the thermal cycler. Fluorescence signal obtained from blank wells (absence of His-SIRT5) was subtracted from each experimental sample. Melting temperature was determined by performing non-linear fitting of normalized thermal denaturation fluorescence signal. All data points after two data points of the maximal raw fluorescence data were deleted, and a “truncated dataset” was generated. The minimal data point in the “truncated dataset” was subtracted from each data point, subsequently each data point was normalized to the maximal data point. The normalized data were then fitted to Boltzmann Sigmoidal curve in GraphPad Prism software (version 5.04).

### RNA extraction and cDNA synthesis (Figure 2b)

1x ice-cold Hank’s balanced salt solution (HBSS, #14175129, Gibco) was added to the human fibroblasts immediately after culture medium was discarded. Fibroblasts were scraped off flasks by scraper, followed by centrifugation at 300 x g, 4°C for 5 min. Cells were washed twice with ice-cold 1x HBSS. Collected cell pellet was then immediately processed to RNA extraction using RNeasy mini kit (#74106, Qiagen), according to the manufacturer’s instructions. Total RNA was dissolved in DNase/RNase-free water, and extracted RNA was kept on ice onwards from now. RNA concentration and quality were measured by a Nanodrop spectrophotometer (IsoGen Life Science). RNA integrity was checked by capillary electrophoresis using Agilent D1000 ScreenTape on the TapeStation 2200 (Agilent Technologies). Subsequently, 1 µg of total RNA was reverse transcribed to cDNA by using an iScript kit (#1708891, Bio-Rad) on a Mastercycler (Eppendorf), with 5 min at 25°C, 30 min at 42°C, and 5 min at 85°C. one cDNA synthesis reaction with an RNA sample without reverse transcriptase (the minRT reaction) was included as a negative control for further real-time RT-PCR analysis. cDNA was stored at -20°C until use.

### Real-Time RT-PCR Analysis (Figure 2b)

Transcript expression was measured using iQ SYBR Green Supermix (#1725006CUST, Bio-Rad), and the fluorescence signal was monitored with a CFX96 Real-Time PCR Detection System. The PCR reaction program was 3 min at 95°C, 40 cycles of 15 s at 95°C and 45 s at 60°C, followed by the melt-curve analysis. Low-level expressed gene ribosomal protein S15 (RPS15) was preamplified with SsoAdvanced PreAmp Supermix (#1725160, Bio-Rad), with 3 min at 95°C, and 12 cycles of 15 s at 95°C and 4 min at 58°C. A negative control (a sample without reverse transcriptase) was included in the preamplification. cDNA from all (preamplified) samples was pooled and serial dilutions were made for standard curves. Samples were diluted 100 fold for PCR reaction, and two negative controls (DNase/RNase-free water and the minRT reaction) were also included in the real-time RT-PCR analysis. The primer sequences are listed in Table 2. SIRT5 gene expression level was normalized to expression level of three reference genes, beta-2 microglobulin (B2M), RPS15, and erbb2 interacting protein (ERBB2IP) in each fibroblast cell line. RNA of each fibroblast cell line was isolated from three independent batches of cultures, and all the sample were analysed in the same RT-PCR run. Duplicates were used for each sample.

### Whole exome sequencing

Whole exome sequencing (WES) and data analysis were performed as described previously^44,45^. In short, exome enrichment was performed using the SureSelect Human All Exon 50 Mb Kit V5 (Agilent). Sequencing was done on a HiSeq4000 (Illumina) with a minimum median coverage of 80×. Read alignment to the human reference genome (GrCH37/hg19) and variant calling was performed at BGI (Copenhagen) using BWA and GATK software, respectively. Variant annotation was performed using a custom designed in-house annotation. Intronic variants (except for splice sites), synonymous changes, and common variants were filtered and excluded from the initial datasets. Patient data were first analyzed using a custom-made virtual gene panel containing mitochondrial disease genes (as described in OMIM) as well all other genes known to encode mitochondrial proteins. As no disease-causing variants were detected, the entire exome was investigated for rare, protein damaging variants. This was done by comparison with the GnomAD dataset, dbSNPv132 and our in-house variant database with MAF depending on mode of inheritance.

### Vertebrate animals study approval

All mice were housed at the Biomedical Science Research Building (University of Michigan). All vertebrate animal experiments were approved by and performed in accordance with the regulations of the University Committee on Use and Care of Animals (University of Michigan).

### Mutagenesis of SIRT5 plasmid (Figure S2c-S2g)

His tagged SIRT5 expression plasmid (SIRT5-pET15b)^46^ was used to generate His tagged SIRT5-P114T expression plasmid (SIRT5^P114T^-pET15b). Site-directed mutagenesis of WT SIRT5 was performed using the QuickChange II-directed mutagenesis kit (#200523, Agilent Technologies), according to the manufacturer’s instruction. The introduction of the mutation (c.340C>A, p.P114T) was confirmed by Sanger sequencing. Mutagenesis primers used are listed in table 2.

### Purification of recombinant proteins (Figure S2c-S2g)

Recombinant His tagged WT SIRT5 and P114T variant SIRT5 proteins were purified using Ni-NTA resin as described previously^46^.

### SIRT5 activity assay (Figure S2e-S2f)

All reactions were performed in 96-well black microplate (Corning No. 3603) with a reaction volume of 50 μL per well. The assay buffer contained 50 mM Tris, 137 mM NaCl, and 2.7 mM KCl at pH 8.0. Reaction wells contained SIRT5 proteins (wild-type or variant, 0.2 μM), fluorescent substrate (Ac-K(Suc)-AMC or Ac-K(Glu)-AMC or Ac-K(Mal)-AMC, 10 μM), NAD^+^ (200 μM), and inhibitors (if applicable) in assay buffer^47^. Control wells without enzyme were incubated in each plate with aim of excluding the interference of fluorescent compounds. The reactions were incubated for 2 h at 37 ℃ and 140 rpm. Subsequently a solution (50 μL) containing 3∼4 U/μl trypsin (Sigma-Aldrich No. T8003) and 8 mM nicotinamide was added, followed by further incubation for 20 min at 37 ℃ and 140 rpm. Fluorescence intensity was then measured using a microplate reader (BioTek Synergy H1, λex = 360 nm, λem = 460 nm, Optics: Top). All experiments were run at least three times independently.

### Purification of recombinant proteins for crystallization and thermal shift assay

The wild-type SIRT5 (WT) gene (residues 32-302) and a variant form (P114T) were each cloned into a His_6_-TEV (Tobacco Etch Virus) expression vector (pMCSG7) and purified by the following protocol. The constructs were transformed into Rosetta^2^ cells, then grown in Terrific Broth and induced with 0.4 mM isopropyl β-D-1-thiogalactopyranoside overnight at 20 °C. The pelleted cells were lysed by sonication in 25 mM Tris-HCl pH 7.0, 200 mM NaCl, and 0.1% 2-mercaptoethanol with protease inhibitors. The supernatant was cleared by centrifugation and incubated with Ni-NTA resin (Qiagen) pre-equilibrated with lysis buffer. The resin was washed with 25 mM Tris-HCl pH 7.0, 200 mM NaCl and 10 mM imidazole, then the protein eluted with 25 mM Tris pH 7.0, 200 mM NaCl, and 200 mM imidazole. The His-tag was removed with TEV protease during dialysis against 25 mM Tris-HCl pH 7.5, 150 mM NaCl, and 1 mM DTT overnight at 4 °C, followed by reapplication to Ni-NTA resin. The cleaved protein was concentrated and applied to a Superdex 75 (GE Healthcare) pre-equilibrated with 20 mM Tris-HCl pH 7.8 and 150 mM NaCl. The purified proteins were concentrated to 12 mg/mL and stored at -80 °C for crystallization studies.

### Crystallization and Structure Determination of WT and P114T SIRT5

Apo SIRT5 WT and apo P114T proteins were crystallized by hanging drop vapor diffusion at 20 °C. Crystals of apo SIRT5 WT grew in drops containing 0.75 µL of protein and 0.75 µL of well solution (0.1 M MES 6.5, 25% PEG 4K). Crystals of apo P114T grew in drops containing 0.5 µL protein and 0.55 µL well solution (0.1 M MES pH 6.5, 20% PEG 10K, 10 mM praseodymium (III) acetate hydrate). SIRT5 WT crystals were cryoprotected in well solution containing an additional 20% ethylene glycol and flash cooled in liquid nitrogen, whereas Apo P144T crystals were cryoprotected in well solution containing 20% glycerol.

Diffraction data were collected on Advanced Photon Source LS-CAT beamlines 21-ID-G (WT and P114T) and then processed using HKL2000^48^. All structures were solved by molecular replacement in Phaser^49^ using a deposited structure of human SIRT5 (PDB ID: 2B4Y) sans ligand as the search model. Iterative rounds of model building and refinement were performed using COOT^37^ and Buster^50^, respectively. Data collection and structural refinement statistics are shown in Table S2.

The structure of the apo form of WT SIRT5 was solved to 2.25 Å resolution in space group P2_1_ with two protein chains per asymmetric unit. Residues 38-302 were modeled in both chains, except for a disordered loop region from 278 to 284 in both chains and 253 to 256 in chain B. The structure of the apo form of P114T was solved to 2.70 Å resolution in space group P2_1_with two protein chains per asymmetric unit. Residues 39-302 were modeled in chain A, except for a disordered loop region from 278 to 284. Residues 37-302 were modeled in chain B, except for two disordered loop regions (278 to 284 and 251 to 253).

### Thermal Shift Assay (Figure S2g)

To determine the melting temperatures for SIRT5 proteins (WT, P114T, and H158Y), 5 uL of buffer (25 mM HEPES 7.0, 200 mM NaCl) containing 2x PTS dye (Applied Biosystems) and 5 uL of 2x protein (5 mg/mL in 25 mM HEPES 7.0, 200 mM NaCl) were added to a 384-well plate (Thermofisher). The plate was centrifuged at 1000 x g for 1 minute. The differential scanning fluorimetry experiment was performed in a Quantstudio7 (Thermofisher) using continuous ramping mode from 25 °C to 95 °C with a scanning rate of 0.03 °C/sec and results analyzed using Protein Thermal Shift software version 1.3 (Applied Biosystems). All assays were performed in quadruplicate.

### Generation of SIRT5 P114T mice

CRISPR/Cas9 technology was used to introduce a Pro114Thr mutation in mouse *Sirt5* exon 4 (Ensembl exon, exon_id=ENSMUSE00000440334.3) on chromosome 13. CRISPR.TEFOR.NET (Haeussler et al., 2016) was used to select single guide RNAs (sgRNAs). Preference for sgRNAs was given to targets that cut near Sirt5 amino acid 114. Two targets predicted to be highly specific and active were tested: sgRNA C181G1 (table 2) with a CFD specificity score of 88 (Doench et al. 2016) and sgRNA C181G2 (table 2) with a CFD specificity score of 89 (Doench et al. 2016). sgRNAs for Cas9 targets were obtained in the form of chemically modified synthetic sgRNA from Synthego.com (Hendel et al., 2015). Recombinant *Streptomyces pyogenes* enhanced specificity Cas9 endonuclease was obtained from MilliporeSigma (ESPCAS9, Slaymaker et al. 2016). C181G1 and C181G2 were assembled into sgRNA/Cas9 ribonucleoprotein (RNP) complexes and tested individually by mouse zygote microinjection to determine if they induced chromosome breaks. Briefly, zygotes were cultured in vitro to the blastocyst stage (about 64 cells per blastocyst) after RNP pronuclear microinjection. DNA was extracted from individual blastocysts and subjected to PCR and amplicon DNA sequencing to identify small insertions/deletions at the Cas9/gRNA cut sites (Sakurai et al., 2014). Primers used to amplify the 5’ sgRNA cut site are listed in table 2. The amplicon size was 641 bp. Amplicons were submitted for Sanger sequencing to determine if the sgRNAs induced by chromosome breaks as shown by “peaks-on-peaks” patterns in Sanger chromatograms that indicate the presence of multiple PCR templates caused by non-homologous endjoining of chromosome breaks. sgRNA C181G1 did not induce chromosome breaks while C181G2 did induce chromosome breaks (Fig. S2a). An Ultramer® single-stranded oligodeoxynucleotide (ssODN, Integrated DNA Technologies) was synthesized to repair chromosome breaks in zygotes through homology directed repair (Jasin and Haber, 2016). This to mutate Pro114 to Pro114Thr, and to block Cas9 cleavage subsequent to repair. The ssODN sequence is listed in table 2. The Pro114Thr amino acid codon change is shown in bold. Three silent codon changes (underlined codons) were made to block Cas9 cleavage after ssODN chromosome break repair.

To generate mice carrying Pro114Thre 309 zygotes obtained from the mating of B6SJLF1 (Jackson Laboratory stock no. 100012) female and male mice were used for pronuclear microinjected as described (Behringer et al. 2014). The microinjection mixture contained sgRNA C181G2 (30 ng/ul) complexed with ESPCAS9 protein (50 ng/µl) and 10 ng/ul ssODN. Surviving zygotes were transferred to pseudopregnant females. 110 potentially gene edited G0 founder pups were screened for the Pro114Thr by PCR amplicon sequencing. Two apparently homozygous G0 founder animals carrying the mutation were mated to wild type mice for germline transmission.

### Genotyping Strategy

To distinguish between WT, *Sirt5* P114T heterozygotes, and homozygotes, PCR was performed using DNA extracted from the tail snips of 21 days old mice. Primers used for PCR amplification are shown in table 2. PCR products (380bp in size) were then digested using AvaI (NEB #R0152L) and separated on agarose gel. AvaI cuts the WT amplicon after bp 85 and produces two fragments that correspond to 85 bp and 295 bp in size. AvaI does not cut P114T amplicon and thus, P114T homozygote amplicon remains as 380 bp in size. AvaI digestion of PCR product from P114T heterozygote results in three bands; 85 bp, 295 bp, and 380 bp.

### Mouse embryonic fibroblasts

P114T heterozygous mice were interbred to generate mouse embryonic fibroblasts (MEFs) from day 13.5 embryos by standard methods. Genotypes were confirmed by Sanger sequencing. Wild-type, heterozygous P114T and homozygous P114T MEFs were cultured in DMEM (Gibco) containing 4.5g/L glucose, 1mM sodium pyruvate, 4mM L-glutamine, 1% non-essential amino acids, 100units/mL penicillin, 100µg/mL streptomycin, 10mM HEPES, 115μM 2-mercaptoethanol, and 15% heat-inactivated FBS, and were grown in a humidified chamber at 37°C containing 5% CO_2_ and 3% O_2_. Where indicated, MEFs were treated 8 μM of proteasomal inhibitor MG132 for 12 hours.

### Immunoblotting

Mice were anesthetized with isoflurane and immediately sacrificed by cervical dislocation. Tissue samples were flash frozen, pulverized, and resuspended in RIPA buffer (50mM Tris-HCL pH7.4, 150mM NaCl, 1% NP-40, 0.5% Na-deoxycholate, 0.1% SDS, 2mM EDTA, 50mM NaF) supplemented with protease and phosphatase inhibitors, 10mM NAM, and 1μM TSA. Samples were sonicated for 60 seconds on ice (where cells were used sonication was performed for 30 seconds), then spun down at 4°C for 10 minutes at 15,000xg, and the supernatant was collected for protein quantification using a DC Protein Assay (Bio-Rad #5000112). Samples were mixed with Laemmli buffer (62.5mM Tris pH6.8, 2% SDS, 10% glycerol, 5% β-mercaptoethanol, 1% bromophenol blue) and boiled for 10’, resolved by SDS-PAGE, and transferred to PVDF membrane overnight at 4°C using a Bio-Rad Criterion system. Membranes were stained with Ponceau S to assess protein loading, and then blocked using 5% milk in TBST (TBS containing 0.1% Tween-20) at room temperature. Primary antibodies were diluted in 5% BSA in TBST and incubated with the blot overnight at 4°C on a rocker. Secondary incubation was performed at room temperature for 1 hour using either mouse or rabbit secondary antibodies (Jackson ImmunoResearch 115-035-062 or 111-035-045) diluted 1:10000 in 5% milk in TBST. For detection, blots were immersed in a western chemiluminescent HRP substrate (Millipore P90720) and imaged using the GE ImageQuant LAS 4000. Primary antibodies used in this study are listed in table 1.

### Isolation of total RNA

Total RNA was extracted from cultured MEFs using Trizol reagent (Life Technologies #15596018). To remove any contaminating genomic DNA, extracted RNA was treated with RNase-free DNase I (Roche #04716728001) for 1 hour at 37°C, according to manufacturer’s instructions. DNase I treated RNA was cleaned up by phenol: chloroform extraction method.

### qRT-PCR

cDNA was generated from DNase I treated total RNA using MultiScribe Reverse Transcriptase (Applied Biosystems High-Capacity cDNA Reverse Transcription Kit with RNase inhibitor # 4374966) per the manufacturer’s instructions. qPCR was performed with PowerUp SYBR Green Master Mix (Applied Biosystems # A25742), in an Applied Biosystems Step One Plus Real-Time PCR Machine. Data acquisition was performed using the manufacturer’s software and data analysis was performed using a ^ΔΔ^CT approach. The primers used for qPCR are listed in table 2.

### Agilent/Seahorse assay

Cellular energy phenotypes, Real-Time ATP production rate, glycolytic capacity, and mitochondrial respiration parameters were measured using the XFe96 Extracellular Flux Analyzer (Seahorse Bioscience, Agilent Technologies, Santa Clara, CA).

The Seahorse Cell Energy Phenotype Test was performed to measure cellular baseline energy phenotype, stressed energy phenotype, and the metabolic potentials. The assay was performed per manufacturer’s instructions. Briefly, 2x10^4^ SIRT5 WT or P114T MEF were plated in complete MEF growth media into each well of a 96-well Seahorse microplate. Cells were then incubated in 5% CO_2_ at 37°C for overnight. Following incubation, cells were washed twice, incubated in a non-CO2 incubator at 37°C, and analyzed in XF assay media (non-buffered DMEM containing 10mM glucose, 2mM L-glutamine, and 1mM sodium pyruvate, pH 7.4) at 37°C, under basal conditions and in response to a mixture of 1.5μM oligomycin + 1μM fluoro-carbonyl cyanide phenylhydrazone (FCCP). Data were analyzed by the Seahorse XF Cell Energy Phenotype Report Generator. The data were normalized to protein content.

The Seahorse XF Real-Time ATP Rate Assay Kit (Agilent) was used to simultaneously measure the basal ATP production rates from mitochondrial respiration and glycolysis. The assay was performed per manufacturer’s instructions. Briefly, 2x10^4^ SIRT5 WT or P114T MEFs were plated in complete MEF growth media into each well of a 96-well Seahorse microplate. Cells were then incubated in 5% CO_2_ at 37°C for overnight. Following incubation, cells were washed twice, incubated in a non-CO2 incubator at 37°C, and analyzed in XF assay media (non-buffered DMEM containing 10mM glucose, 2mM L-glutamine, and 1mM sodium pyruvate, pH 7.4) at 37°C, under basal conditions and in response to 1.5μM oligomycin, and 0.5μM rotenone/0.5μM antimycin A. Data were analyzed by the Seahorse XF Real-Time ATP Rate Assay Report Generator. ATP production rates were normalized to protein content.

To measure glucose- or galactose-dependent mitochondrial respiration, mitochondrial stress tests were performed per manufacturer’s instructions. Briefly, 2x10^4^ SIRT5 WT or P114T MEFs were plated in complete MEF growth media into each well of a 96-well Seahorse microplate. Cells were then incubated in 5% CO_2_ at 37°C for overnight. Following incubation, cells were washed twice, incubated in non-CO_2_ incubator at 37°C, and analyzed in XF assay media (non-buffered DMEM containing 25mM glucose or 25mM galactose, 2mM L-glutamine, and 1mM sodium pyruvate, pH 7.4) at 37°C, under basal conditions and in response to 1.5μM oligomycin (Sigma), 1μM fluoro-carbonyl cyanide phenylhydrazone (FCCP) (Sigma) and 0.5μM rotenone (Sigma)/0.5μM antimycin A (Sigma). Data were analyzed by the Seahorse XF Cell Mito Stress Test Report Generator. Oxygen consumption rate (OCR) (pmol O_2_/min) values were normalized to the protein content.

Glycolytic parameters were measured by performing glycolysis stress tests according to manufacturer’s instructions. Briefly, 2x10^4^ SIRT5 WT or P114T MEFs were plated in complete MEF growth media into each well of a 96-well Seahorse microplate. Cells were then incubated in 5% CO_2_ at 37°C for overnight. Following incubation, cells were washed twice, incubated in a non-CO_2_ incubator at 37°C, and analyzed in XF assay media (non-buffered DMEM containing 2mM L-glutamine, pH 7.4) at 37°C, under basal conditions and in response to 10mM glucose (Sigma), 1.5μM oligomycin (Sigma), and 50mM 2-deoxy-D-glucose (Sigma). Data were analyzed by the Seahorse XF Cell Glycolysis Stress Test Report Generator. Extracellular acidification rate (ECAR) (mpH/min) values were normalized to protein content.

### Brain immunofluorescence

To prepare brain tissues for immunofluorescence, mice were perfused with saline and 4% PFA and tissue was removed. Brains were immersed in 30% sucrose for 24 hours and then fixed in 4% PFA overnight at 4°C. Fixed tissues were embedded in OCT and frozen. Frozen tissues were cut on a Leica CM1900 cryostat into 10 μm sections and placed on Thermo Scientific Superfrost Plus microscope slides. For staining, slides were incubated in a solution with 0.1% Triton, 10% goat serum, and 1% BSA in PBS for 20 minutes. Slides were placed in blocking solution (10% goat serum, 1% BSA in PBS) for 40 minutes before incubating in primary antibody (1:500 NeuN, 1:500 GFAP, 1:250 Iba1) diluted in blocking solution overnight at 4°C. Slides were washed 3 times in PBS and incubated with secondary antibody (1:500) diluted in blocking solution for 1 hour at room temperature. Slides were then washed 3 times in PBS and mounted with Vectashield plus DAPI (Vector Laboratories). 40x images of midline sagittal sections were captured on a Nikon A-1 confocal microscope. Images were captured from 3 different brain regions: cerebellum (lobules 3 and 5), hippocampus (CA2), and cortex. 2 Images were captured/brain region/mouse. Staining intensity for each image was measured using ImageJ. Counts of GFAP or Iba1 positive cells were done using CellProfiler. Intensity and count values from the 2 images from each brain region were averaged for each sample. Significance was determined using GraphPad Prism.

The primary and secondary antibodies used in this study are listed in table 1.

### Statistics

Statistical analysis were conducted using Graphpad Prism. Multiple-group comparisons were analyzed using two-way ANOVA followed by Sidak’s correction for multiple comparisons. For analysis of two groups, Welch’s t-test was used. A p-valu*e* of less than 0.05 was considered statistically significant. Scatter plots include median and interquartile range markers. Significance markers: (*) p < 0.05, (**) p < 0.01, (***) p < 0.001, (****) p < 0.0001.

## Data Availability

The primary data that support the findings of this study are available from the corresponding authors upon reasonable request.

## Acknowledgments

The authors would like to thank the University of Michigan Transgenic Animal Model Core for generation of the SIRT5 P114T mouse strain, and members of the Lombard Lab for useful feedback. This project was supported by R01GM101171 (DL) and R01CA253986 (DL/NN). TY has received financial support from the China Scholarship Council (Grant no. 201606350170). Use of the Advanced Photon Source was supported by the U. S. Department of Energy, Office of Science, Office of Basic Energy Sciences, under Contract No. DE-AC02-06CH11357. Use of the LS-CAT Sector 21 was supported by the Michigan Economic Development Corporation and the Michigan Technology Tri-Corridor for the support of this research program (Grant 085P1000817).

## Author contributions

Project conception: TY, SK, APL, JS, NN, RR, JNS, VDB, DBL; performed experiments: TY, SK, MES, RVJ, MA, JZ, TJK, EMP, TS, JS; contributed reagents and/or patient data: FA, AM, RR; project management: JK, VDB, DL

## Supplementary Figure Legends

**Figure S1:**
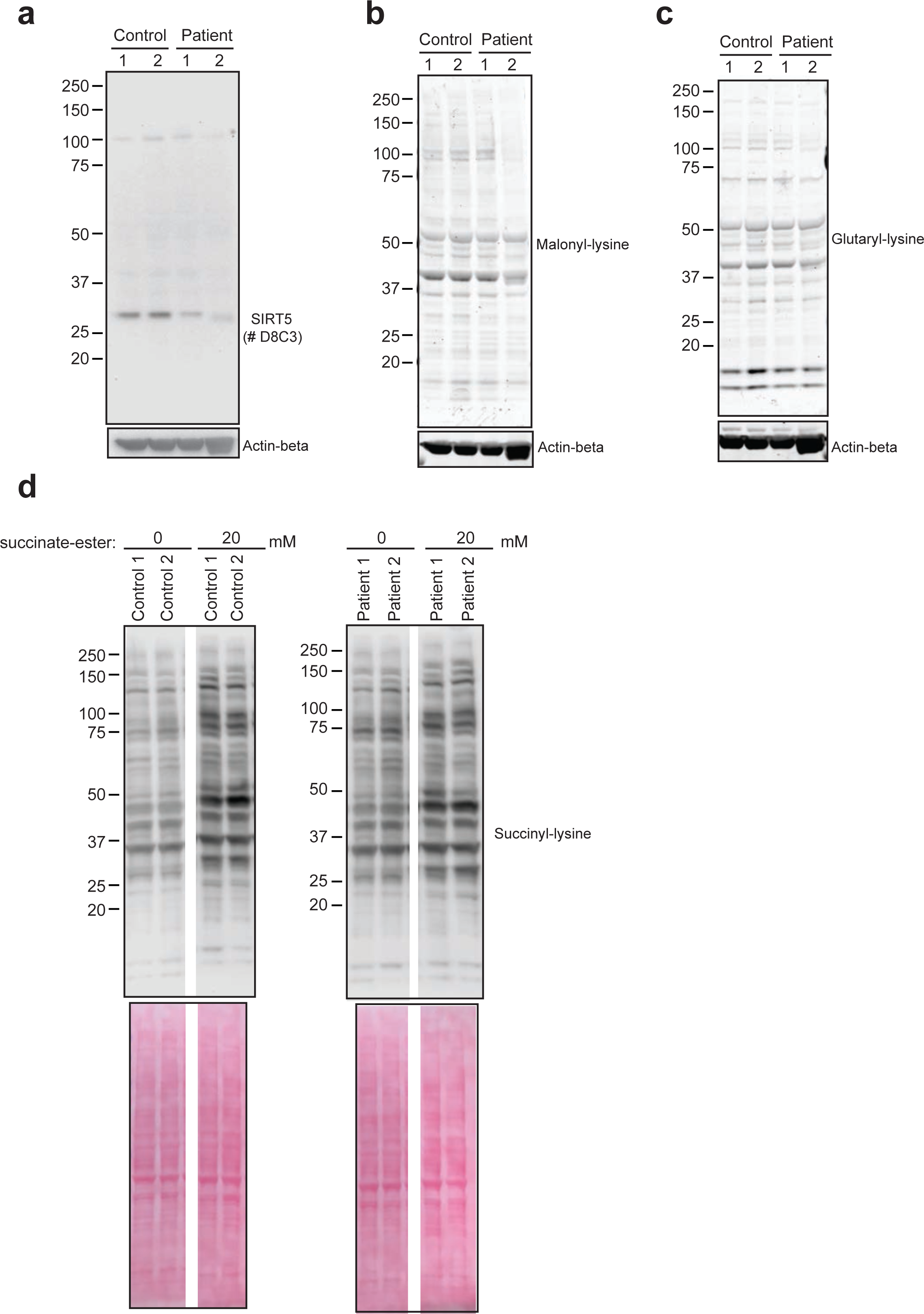
(a) Immunoblot for SIRT5 using different anti-SIRT5 antibody in whole cell lysates from control and patient fibroblasts. Beta-actin served as loading control. (b-c) Immunoblots for malonyl-lysine and glutaryl-lysine with loading control beta-actin in whole cell lysates from control and patient fibroblasts. (d) Immunoblots showing increased succinylation levels in both control and patient fibroblasts upon exposure to dimethyl succinate-ester, a cell permeable succinate analogue.

**Figure S2:**
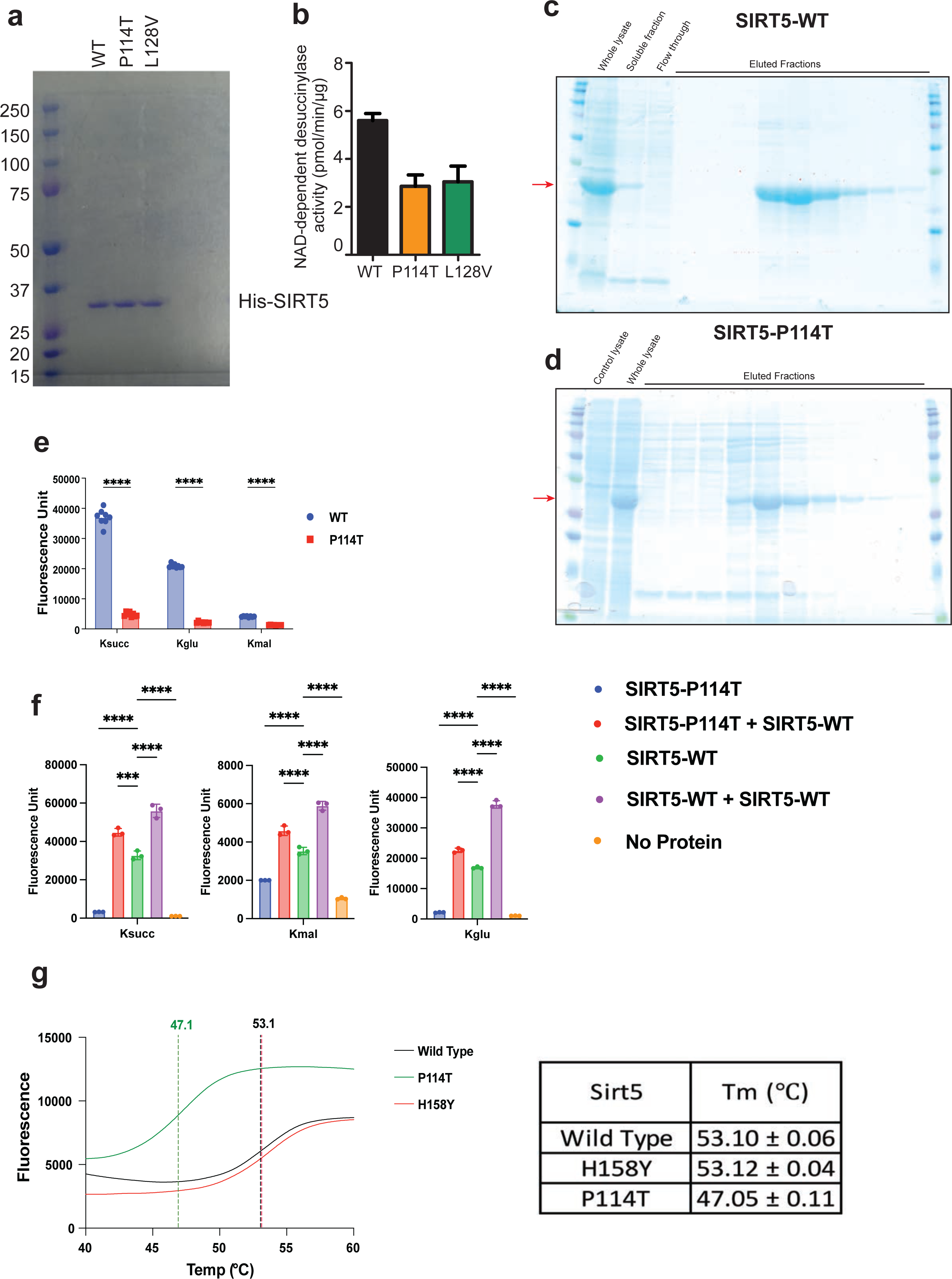
Purification and activity of recombinant WT SIRT5 and SIRT5 P114T variant. (a) SDS-PAGE image of purified recombinant WT SIRT5, SIRT5 P114T, and SIRT5 L128V proteins. (b) NAD dependent desuccinylase activity of recombinant WT SIRT5, SIRT5 P114T, and SIRT5 L128V proteins. (c-d) SDS-PAGE images of purified recombinant WT SIRT5 and SIRT5 P114T proteins. (e) SIRT5 activity assay; the P114T variant displays reduced biochemical activity against all three target marks when compared with WT SIRT5. (g) Thermal shift assay; graph and table show the Tm for the SIRT5 WT, SIRT5 H158Y and SIRT5 P114T variant proteins. Values represent the mean ± SD of four independent experiments.

**Figure S3:**
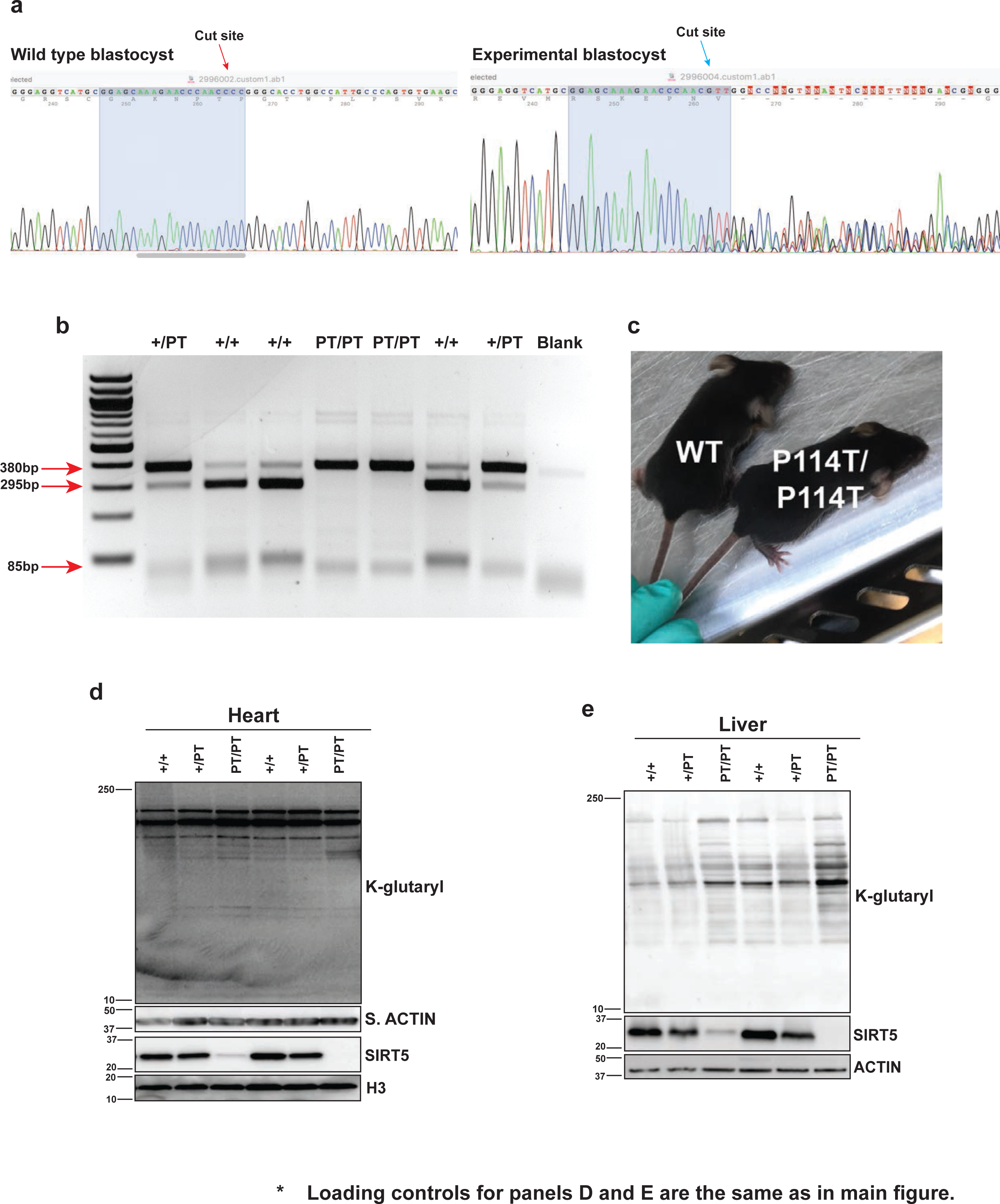
(a) Sanger sequencing peaks showing the CRISPR/Cas9 target site. A DNA fragment spanning the expected Cas9 cut site (red arrows, WT blastocyst, blue arrow, sample blastocyst) was PCR amplified and the product was sequenced. Test blastocyst shows the expected peaks-on-peaks pattern in a chimeric sample. (b) Agarose gel image of genotyping PCR to distinguish between WT, *Sirt5* P114T heterozygotes, and homozygotes. (c) WT and *Sirt5* P114T homozygotes show no apparent morphologic anomalies. (d-e) Immunoblots showing reduced levels of SIRT5 protein and slightly increased levels of Kglu in P114T homozygous tissues (hearts and liver) compared to WT controls.

**Figure S4:**
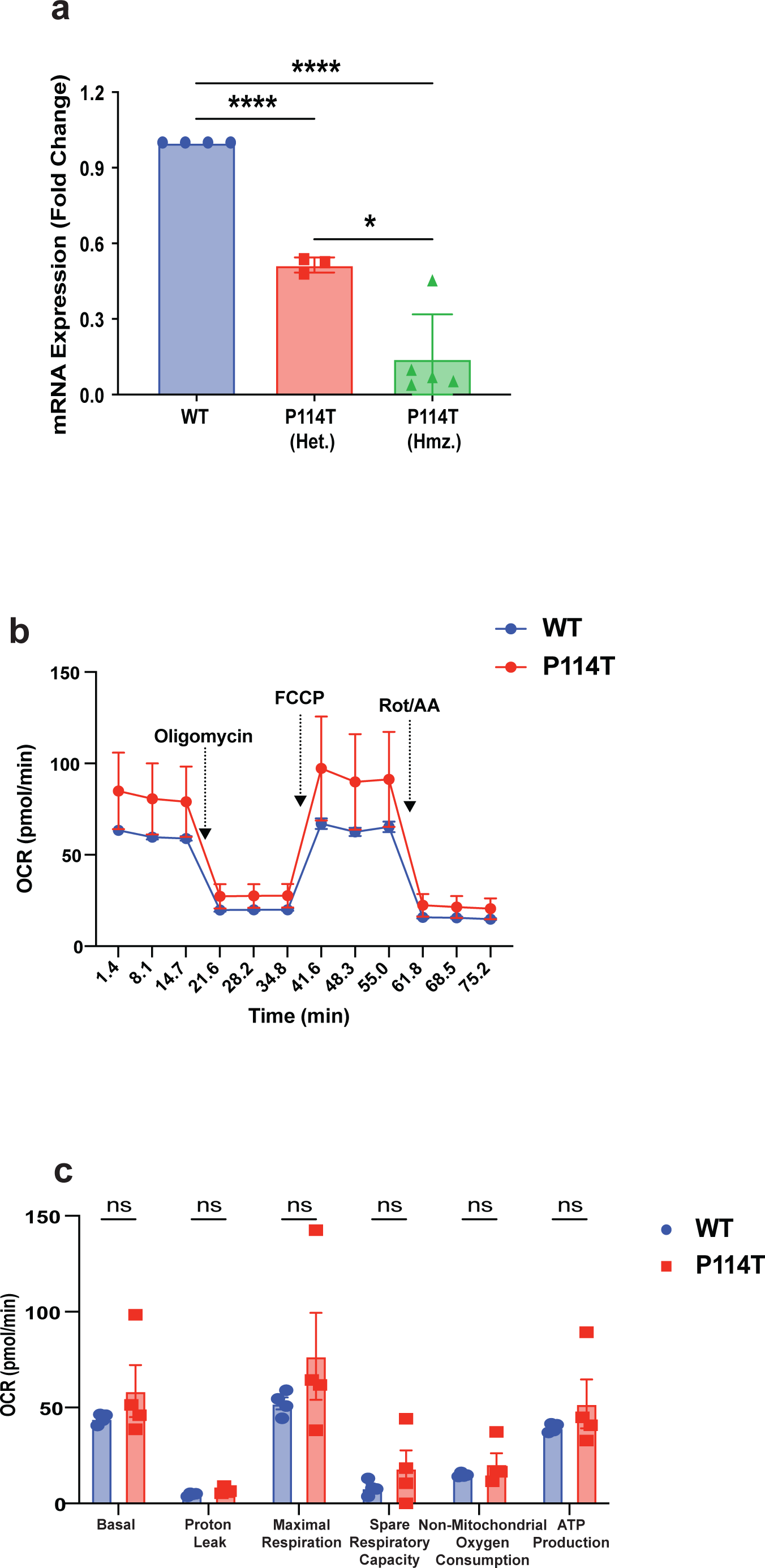
(a) qRT-PCR showing reduced levels of *Sirt5* mRNA in P114T heterozygous and homozygous MEFs compared with WT controls. (b-c) P114T variant MEFs maintain galactose dependent mitochondrial respiration.

**Figure S5:**
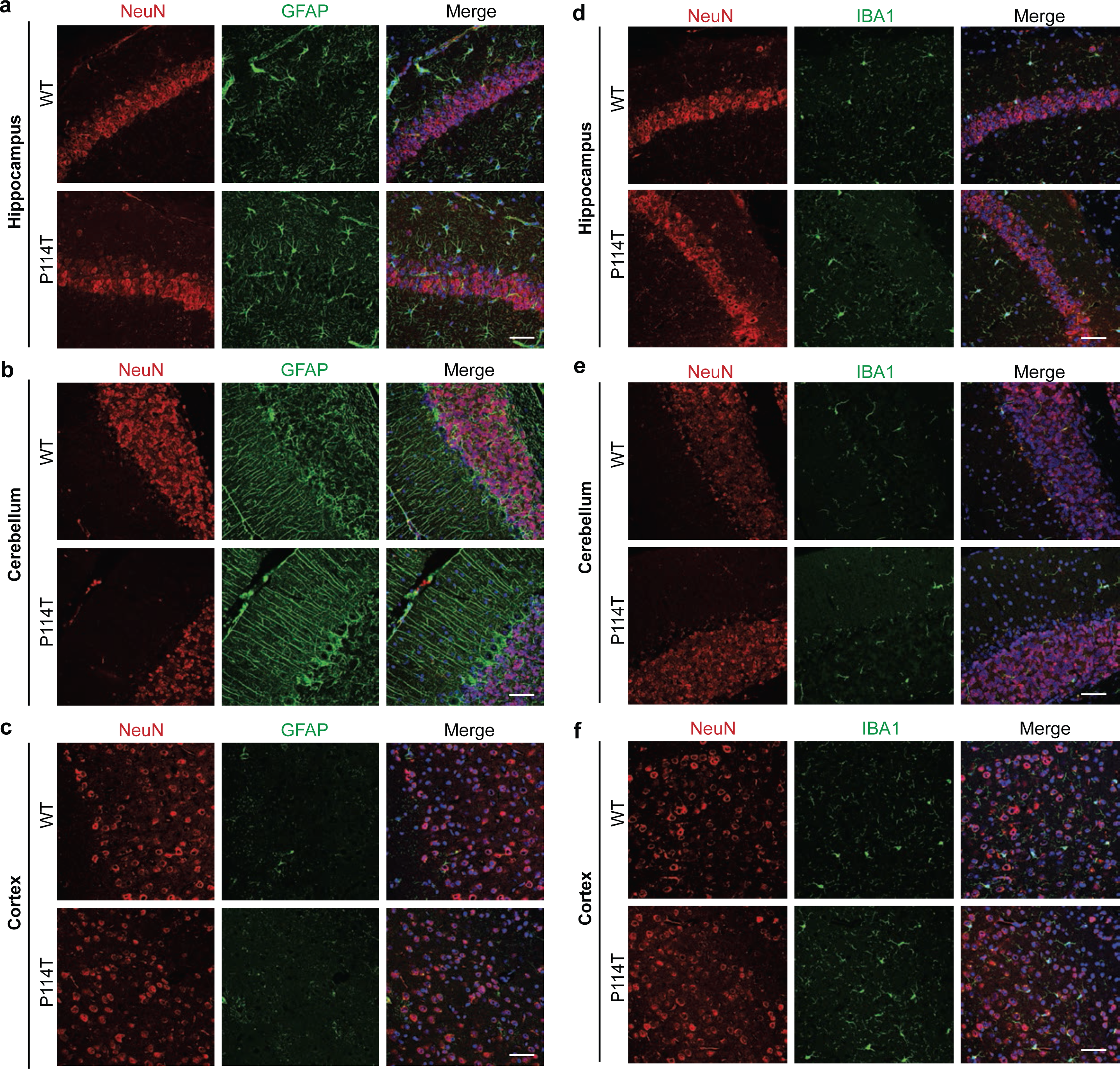
GFAP and Iba1 staining in brains from WT and P114T mice. Brains were collected from adult WT and P114T mice. (a-c) Images show GFAP and NeuN staining in the hippocampus (CA2), frontal cortex, and cerebellum (lobule 3) from WT and P114T mice. (d-f) Images show Iba1 and NeuN staining in the hippocampus (CA2), frontal cortex, and cerebellum (lobule 3) from SIRT5-WT and SIRT5-P114T mice.

**Supplementary Table S1.**

| Enzyme activities (human fibroblasts) | Activity |  | Control range | Unit |
| --- | --- | --- | --- | --- |
|  | Patient 1 | Patient 2 |  |  |
| Complex I | 565 | 770 | 279 - 1076 | mU/UCOX |
| Complex II | 978 | 956 | 375 - 2692 | mU/UCOX |
| Complex III | 1576 | 1327 | 623 - 3534 | mU/UCOX |
| Complex II + III (succ.: cyt.c oxidoreductase; SCC) | 565 | 522 | 269 - 781 | mU/UCOX |
| Complex IV | 377 | 541 | 288 - 954 | mU/UCS |
| Complex V (ATPase) | 1261 | 1434 | 480 - 2705 | mU/UCOX |
| Citrate synthase | 428 | 435 | 151 - 499 | mU/mg |

| Enzyme activities (skeletal muscle) | Activity |  | Control range | Unit |
| --- | --- | --- | --- | --- |
|  | Patient 1 | Patient 2 |  |  |
| Complex I | ND | 107 | 68 - 230 | mU/UCS |
| Complex II | ND | 216 | 76 - 281 | mU/UCS |
| Complex III | ND | 244 | 182 - 1421 | mU/UCS |
| Complex II + III (succ.: cyt.c oxidoreductase; SCC) | ND | 56 | 30 - 263 | mU/UCS |
| Complex IV | ND | 229 | 228 - 1032 | mU/UCS |
| Citrate synthase | ND | 299 | 111 - 604 | mU/mg |

**Table S2:** Crystallography Data Collection and Refinement Statistics.

| Data Collection | Apo P114T | SIRT5 WT |
| --- | --- | --- |
| PDBID | 8GBN | 8GBL |
| Space Group | P2<sub>1</sub> | P2<sub>1</sub> |
| Unit Cell (Å) | a = 41.02<br>b = 114.90<br>c = 56.27<br>$\alpha = 90^\circ$,<br>$\beta = 90.14^\circ$ | a = 41.62<br>b = 112.53<br>c = 56.08<br>$\alpha = 90^\circ$,<br>$\beta = 90.03^\circ$ |
| Wavelength (Å) | 0.9786 | 0.9786 |
| Resolution (Å)<sup>1</sup> | 2.70 (2.75-2.70) | 2.25 (2.25-2.29) |
| Rsym<sup>2</sup> | 0.082 (0.612) | 0.081 (0.756) |
| $\langle I/\sigma I \rangle^3$ | 19.29 (2.64) | 23.91 (3) |
| Completeness (%)<sup>4</sup> | 94.3 (94.2) | 100 (100) |
| Redundancy | 5.4 (5.4) | 7.6 (7.6) |
| **Refinement** |  |  |
| Resolution (Å) | 2.7 | 2.24 |
| R-Factor<sup>5</sup> | 0.1819 | 0.2151 |
| Rfree<sup>6</sup> | 0.2311 | 0.2563 |
| Protein atoms | 3802 | 3791 |
| Ligands | 0 | 0 |
| Water Molecules | 92 | 116 |
| Unique Reflections | 13302 | 23941 |
| R.m.s.d.<sup>7</sup> |  |  |
| Bonds | 0.01 | 0.01 |
| Angles | 1.09 | 1.06 |
| MolProbity Score<sup>8</sup> | 1.65 | 0.91 |
| Clash Score<sup>8</sup> | 3.76 | 1.35 |

<sup>1</sup>Statistics for highest resolution bin of reflections in parentheses.
<sup>2</sup> $R_{\text{sym}} = \frac{\sum_h \sum_j |I_{hj} - \langle I_h \rangle|}{\sum_h \sum_j I_{hj}}$ , where $I_{hj}$ is the intensity of observation $j$ of reflection $h$ and $\langle I_h \rangle$ is the mean intensity for multiply recorded reflections.
<sup>3</sup>Intensity signal-to-noise ratio.
<sup>4</sup>Completeness of the unique diffraction data.
<sup>5</sup> $R\text{-factor} = \frac{\sum_h |F_o| - |F_c|}{\sum_h |F_o|}$ , where $F_o$ and $F_c$ are the observed and calculated structure factor amplitudes for reflection $h$ .
<sup>6</sup> $R_{\text{free}}$ is calculated against a 5% random sampling of the reflections that were removed before structure refinement.
<sup>7</sup>Root mean square deviation of bond lengths and bond angles.
<sup>8</sup>Chen et al. (2010) MolProbity: all-atom structure validation for macromolecular crystallography. Acta Crystallographica D66:12-21.

